# Gut microbial diversity and inferred capacity to produce short-chain fatty acids are tied to acute stress reactivity in healthy adults

**DOI:** 10.1101/2025.06.24.661294

**Authors:** Thomas Karner, Paul A.G. Forbes, David Berry, Isabella C. Wagner

## Abstract

Acute stress triggers the release of stress hormones such as cortisol, increasing stress reactivity and aiding post-stress recovery. Rodent studies revealed that stress reactivity is modulated by the gut microbiota, and few interventional studies have provided evidence for an effect on human cortisol dynamics. However, it remains unclear whether stress reactivity is related to interindividual variations in gut microbial composition and to one’s capacity to produce microbial metabolites such as short-chain fatty acids (SCFAs). To close this gap, we analyzed data from 74 healthy human adults who completed the study in the laboratory and were either exposed to a well-established, standardized intervention that induced acute stress or to a non-stressful control condition (*n* = 35/39 per stress/control group). Stool samples were obtained at baseline, and the gut microbiota were characterized through 16S rRNA gene amplicon sequencing. Cortisol changes were assessed from repeated saliva sampling, paralleled by measurements of subjectively experienced stress. We found that higher gut microbial alpha diversity was associated with higher cortisol and subjective stress reactivity across individuals of the stress group, but not in controls. Cortisol stress reactivity was also associated with the relative abundance of bacterial taxa inferred to encode metabolic pathways for the production of butyrate and propionate, two key SCFAs. The results are the first to highlight the link between gut microbial diversity, inferred SCFA production capacity, and the acute stress response in healthy adults, underscoring the microbiota’s potential to flexibly modulate human psychophysiology in the aftermath of stress.

## 1. Introduction

Acute stress describes the allostatic processes that cause a deviation from bodily homeostasis in response to a perceived threat or challenge (McEwen, 2017; Sapolsky, 2019). On a physiological level, stress is regulated by the hypothalamic-pituitary-adrenal (HPA) axis, which titrates the release of stress hormones such as cortisol (De Kloet et al., 2005). Encountering an acute stressor initiates a rise in cortisol (typically peaking after approx. 20 minutes; Dickerson & Kemeny, 2004; Linz et al., 2019), after which it returns to baseline (approx. 90 minutes post-stressor; Kirschbaum & Hellhammer, 1994; Linz et al., 2019). An exaggerated rise in cortisol and/or a prolonged post-stress recovery after acute stress may occur in individuals who experienced chronic early-life stress or who show heightened threat sensitivity (Heim et al., 2001; Miller et al., 2007), whereas blunted responses may appear in anxiety-related disorders and in depression (Burke et al., 2005; Fiksdal et al., 2019; McEwen, 2017; Miller et al., 2007; Zorn et al., 2017). Both extremes stand in contrast to moderate and well-regulated cortisol dynamics, indicating that time-limited cortisol stress reactivity and post-stress recovery are important markers of a flexible, adaptive stress response (Kudielka et al., 2009; McEwen, 2017; Miller et al., 2007).

Emerging evidence suggests that the microorganisms that inhabit the gastrointestinal tract, collectively termed the gut microbiota, play a crucial role in modulating the stress response (Cryan et al., 2019; Foster et al., 2017). For instance, Sudo and colleagues (2004) found that germ-free mice, which lack a gut microbiota exhibited exaggerated stress reactivity that could be normalized through colonization with *Bifidobacterium infantis,* a probiotic bacterium known to confer health benefits to the host. Gut microbiota composition can be described through diversity (which accounts for the number and/or relative abundance of genetically distinct bacterial taxa). While the overall gut microbiota composition appears highly specific to an individual (Yatsunenko et al., 2012), patients with major depressive disorder were shown to exhibit altered gut microbiota composition (e.g., decreased bacterial diversity compared to healthy controls). Moreover, transplantation of their fecal material to microbiota-depleted rats induced depression-like behavior in these animals (Kelly et al., 2016). In humans, probiotic (thus, beneficial) gut bacteria were shown to buffer against the detrimental effects of acute stress on cognition (Papalini et al., 2019), negative emotions (Steenbergen et al., 2015), to reduce perceived stress, improve sleep quality, and impact cortisol dynamics following acute stress (Boehme et al., 2023). Whether gut microbiota composition is linked to cortisol stress reactivity and post-stress recovery in healthy adults is, however, unclear.

One pathway of how the gut microbiota can affect stress-related neurophysiology is through the production of metabolites such as short-chain fatty acids (SCFAs; Dalile et al., 2019, 2020; O’Riordan et al., 2022). These are mainly derived from the fermentation of dietary (Casterline et al., 1997; Slavin, 2013) and prebiotic fibers (Davani-Davari et al., 2019; Rastall & Gibson, 2015), are produced by probiotic bacteria (Grimaldi et al., 2017), and are central to maintaining gut health, metabolism, and immune function (Wenzel et al., 2020). Supplementation with a mixture of the key SCFAs acetate, propionate, and butyrate (Van De Wouw et al., 2018), or only with butyrate (Wang et al., 2023) was shown to decrease corticosterone release following acute stress induction in chronically stressed mice and in an autism rat model, respectively. In humans, Dalile and colleagues (2020) found that supplementation with the abovementioned SCFA mixture altered SCFA serum levels and attenuated cortisol release after psychosocial stress induction (Dalile et al., 2020; but note that a follow-up study that supplemented only butyrate via colonic delivery did not replicate this effect and found no significant modulation of cortisol dynamics; Dalile et al., 2024). Moreover, the administration of prebiotics, which are known to increase SCFA-producing bacteria as well as SCFA concentrations (Davani-Davari et al., 2019; Grimaldi et al., 2017) was shown to reduce the cortisol awakening response and partly decreased attentional vigilance to negative information (Schmidt et al., 2015). Nevertheless, a clear understanding of whether and to what degree an individual’s gut microbial capacity to produce SCFAs modulates cortisol stress reactivity and post-stress recovery is missing. Addressing this gap could help identify microbial targets to enhance stress resilience, potentially informing dietary or microbiome-based interventions.

In the present study, we analyzed data from 74 healthy human participants tested in the laboratory. Each participant donated a stool sample for characterization of the gut microbiota in terms of overall composition (using 16S rRNA gene amplicon sequencing) and inferred capacity to produce SCFAs. They were then randomly assigned to a stress or control group and underwent an acute stress induction procedure or a non-stressful control intervention, respectively (using a modified version of the Montreal Imaging Stress Task, MIST; Dedovic et al., 2005). Cortisol stress reactivity and post-stress recovery were quantified by repeated salivary cortisol sampling, paralleled by assessments of participants’ subjective stress experience (**Figure 1A**). We hypothesized that cortisol stress reactivity and post-stress recovery would be linked to gut microbiota composition across participants. Specifically, we expected that higher gut microbiota diversity would be associated with lower cortisol stress reactivity and faster post-stress recovery. Because the gut microbiota was shown to impact stress-related neurophysiology via metabolite production of SCFAs (Van De Wouw et al., 2018), we expected that higher gut microbial capacity to produce SCFAs would be coupled with lower cortisol stress reactivity and faster post-stress recovery across participants. Lastly, since both the acute stress response (Adjei et al., 2018; Stefanaki et al., 2022) and gut microbiota composition (Stefanaki et al., 2022) depend on biological sex, we explored whether the obtained results differed between biological males and females.

**Figure 1.**
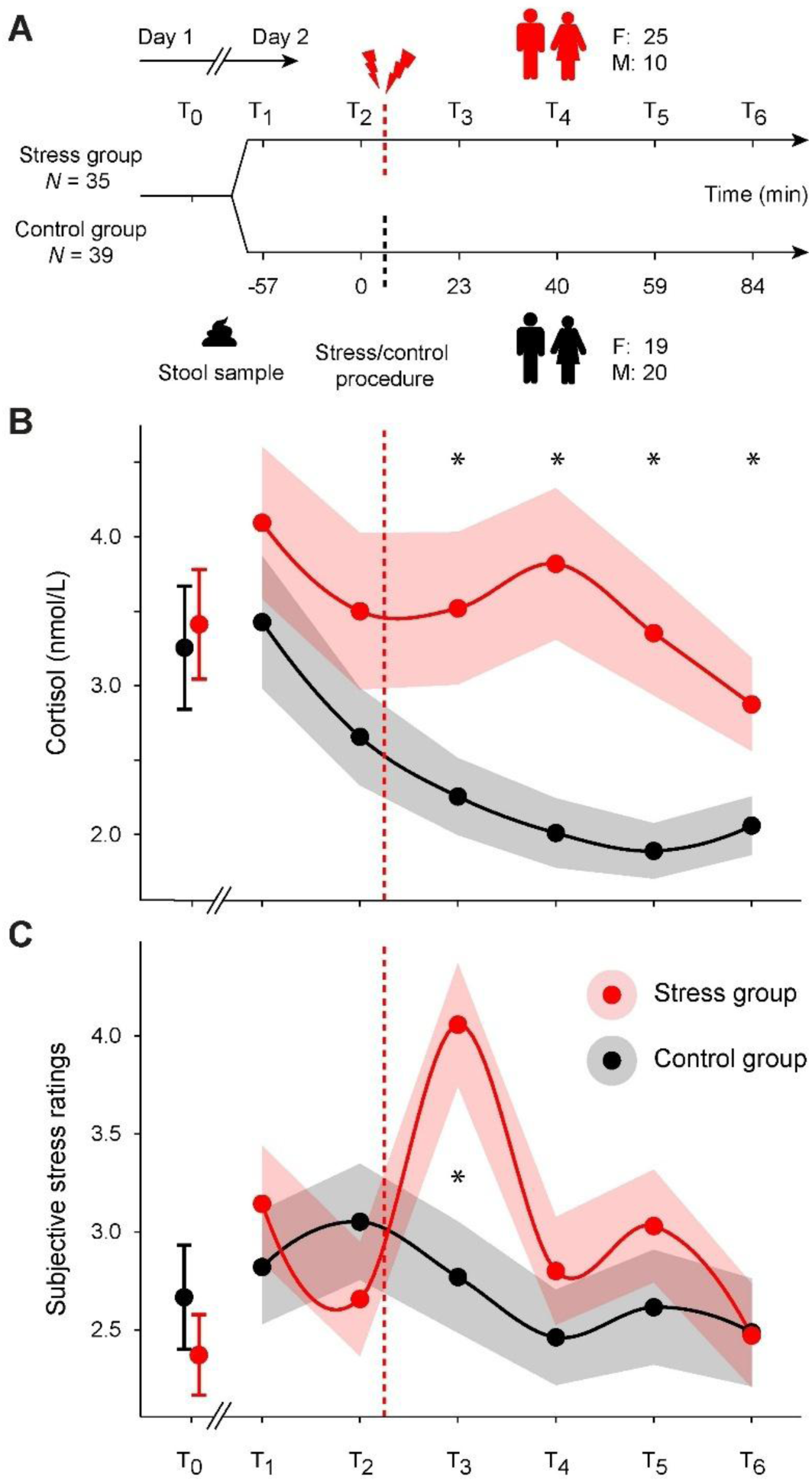
Study timeline, cortisol dynamics, and subjectively experienced stress over time. **(A)** The stress and control groups were tested on two separate study days, approx. one week apart (F/M indicates the number of biological females/males). Stool samples were produced at baseline (i.e., either on the evening of day 1, or prior to the start of the experimental session on day 2, see Methods). All sessions (day 1, 2) took place in the afternoon and sessions timings were kept constant within participants (i.e., starting either at ∼14:15 p.m. or at ∼17:15 p.m., see Methods). Timepoints (T) indicate the collection of salivary cortisol and ratings of subjectively experienced stress. Dashed lines indicate the timing of the acute stress intervention (red) or the control condition (black). Time (minutes) is displayed relative to the study baseline (T2), which directly preceded the onset of the acute stress intervention. Note that a stress reminder was also given after T4 (not shown here; see Methods). **(B)** Salivary cortisol concentrations (nmol/L; statistical comparisons were performed using the log-transformed values), and **(C)** ratings of subjectively experienced stress over time. Values represent the respective group mean, shaded areas represent the standard error of the mean (s.e.m.). * denotes significant group differences (*p* < 0.05).

## 2. Methods

### 2.1. Participants

Eighty healthy participants were recruited through the University of Vienna’s online recruitment platform and were invited to the laboratory (62.5% females, age range: 18-34 years). Participants had no history of neurological or psychiatric diagnosis and did not report any gastrointestinal diseases or symptoms thereof (such as prolonged diarrhea or constipation within the last 3 months). The use of antibiotics, probiotics, or other medications (e.g., steroids, cytokines, proton pump inhibitors) within the past three months was an exclusion criterion since these substances were shown to influence the gut microbiota (Palleja et al., 2018; Vich Vila et al., 2020; Zmora et al., 2018). Also, the intake of hormonal contraceptives was an exclusion criterion, since they were shown to blunt stress hormone levels in females (Kirschbaum et al., 1999; Nielsen et al., 2013). The majority of participants were University students who received course credits or monetary compensation. Four participants were excluded from the analyses because they did not understand the experimental instructions and/or did not engage with the task, one participant was excluded due to abnormal cortisol values (showing extremely low values and no variability across timepoints, likely indicating a sampling error, range: 1.13-2.49 nmol/l, mean: 1.5 nmol/l), and one was excluded due to technical difficulties with the experimental equipment. This resulted in a final sample size of 74 participants who were randomly assigned to the stress group (*n*_stress_ = 35, 25/10 females/males, 71.4/28.6%, age range: 18-34 years, mean age: 22 years) or the control group (*n*_control_ = 39, 19/20 females/males, 48.7/51.3%, age range: 18-33 years, mean age: 23 years). Although the stress group contained relatively more biological females, the distribution of biological sexes did not significantly differ between the groups (Chi-squared test, χ² = 3.060, df = 1, *p* = 0.080). All participants provided written informed consent prior to entering the study. The study procedures were conducted in accordance with the Declaration of Helsinki (World Medical Association, 2013) and were approved by the ethics committee of the University of Vienna, Austria.

### 2.2. Study design

The study took place across two consecutive study days (**Figure 1A**). For measurements of salivary cortisol, participants were instructed to refrain from food and caffeine intake (they were asked to only drink water, but no caffeinated drinks), from brushing their teeth, and from chewing gum up to 2 hours before the start of the experimental sessions on both days. They were also instructed to refrain from physical activity (sports), cigarette smoking, alcohol consumption, and consumption of drugs or medications for up to 24 hours before the start of the first experimental session on day 1 until the end of their study participation on day 2.

On day 1, participants completed initial ratings (T_0_) of their subjective stress experience and their mood (“In this moment I am feeling…?”, items: “stress”, “happy”, “bad”, “anxious”, “disgust”, “sad”, “calm”, “surprise”, “angry”, response via a 7-point Likert scale ranging from “not at all” to “very”). They provided a saliva sample (∼20 minutes after arrival at the laboratory; note that a T_0_ cortisol sample was missing for one participant in the control group, and we interpolated the missing value), and were instructed on how to perform the stool sampling (see below for details on the sampling procedures).

On day 2, participants received the task instructions depending on whether they belonged to the stress or control group. Since this study was part of a larger project that tested the effects of acute stress on the neural underpinnings of spatial navigation, participants then completed a (non-stressful) object-location task (Doeller et al., 2010) within the MR scanner that consisted of four separate task runs (∼15 minutes each, not further discussed here). The acute stress induction took place ∼30 minutes after the start of the MRI measurements (i.e., after 2 task runs of the object-location task) inside the MR scanner (see below for details). After ∼45 minutes (i.e., after the third task run of the object-location task), another acute stress induction took place that acted as a “stress reminder”. This was done to ensure that both remaining runs of the object-location task would be impacted equally by the acute stress induction (Forbes et al., 2024; Vogel et al., 2015). Overall, participants’ subjective stress experience and salivary cortisol were assessed at six time points on day 2 (approx. minutes pre/post main stress induction; T_1_:-57 min; T_2_: 0 min (stress induction); T_3_, +23 min; T_4_, +40 min; T_5_, +59 min; T_6_, +84 min). To account for inter-individual differences in trait anxiety and perceived stress, participants further completed online versions of the General Anxiety Disorder-7 (GAD-7; Spitzer et al., 2006) and the Perceived Stress Scale (PSS; Cohen, 1988).

### 2.3. Acute stress induction procedure

On day 2, participants of the stress group were exposed to a modified version of the Montreal Imaging Stress Task (MIST) that combined an arithmetic challenge with social evaluation (Dedovic et al., 2005; Forbes et al., 2024). At the beginning of the session, participants were told the cover story, which was that the study revolved around the link between arithmetic performance and cognition and that it was important that they performed well on the arithmetic challenge. They were told that their performance would be video-recorded and evaluated. The participants then completed two task runs of the unrelated object-location task, after which the main acute stress intervention took place. This comprised two runs of the computerized arithmetic challenge (6 minutes each), during which participants solved arithmetic problems of increasing difficulty under time pressure. They were shown a live stream via webcam of the experimenter taking notes. Following a standardized feedback protocol, the experimenter gave negative feedback on participants’ performance and told them that their data was unusable if their performance was too low compared to the alleged group average. At certain times, the experimenter interrupted the arithmetic challenge and told participants that the task needed to be started again due to their insufficient performance. The experimenters were wearing white lab coats throughout the session.

After the arithmetic challenge, participants completed a counting task (3 minutes) that was adapted from the Trier Social Stress Test (TSST; Kirschbaum et al., 1993). The experimenter entered the MRI scanner room, stood next to the scanner bed, and instructed the participant to count down from 2043 in steps of 17. They were instructed to start over when making a mistake and to respond faster, maximizing pressure through social evaluation. Experimenters also attempted to distract them from counting by urging them to respond faster or speak more loudly while they were calculating, increasing the likelihood of errors. The experimental session commenced with the third run of the object-location task and another (shorter) acute stress intervention that only included the arithmetic counting task, but with different numbers.

The control group participants were told the same cover story but underwent a non-stressful version of the MIST. They were not told that their performance would be video recorded and evaluated. While the control group still completed the arithmetic challenge with matched difficulty and overall duration, no video live stream of the experimenter was shown, and no negative feedback was provided. The experimenters also did not wear white lab coats during the session. The instructions for the counting task were given via the MRI scanner intercom rather than in person, and participants were counting quietly in their heads from 0 in steps of 5, and, after 3 minutes, reported the number they had reached. The counting task was repeated after the third run of the object-location task. After the study completion, all deception was resolved, and participants were told about the actual study goal.

### 2.4. Salivary cortisol sampling and analysis

For salivary cortisol sampling, participants were instructed to keep the cotton pad in their mouth for 60 seconds, without chewing (Salivette, Sarstedt, Germany). The collected samples were refrigerated for the duration of the experimental session and were subsequently transferred to freezer storage (-25 °C). Upon completion of data collection for the entire study, samples were shipped to the laboratory of Prof. Kirschbaum (Technical University Dresden, Germany) and were analyzed using a commercially available chemiluminescence immunoassay with high sensitivity (IBL International, Hamburg, Germany). Of the 560 samples collected, 537 were analyzed in duplicates, 16 were analyzed as singlets due to limited saliva volume, and 7 were excluded due to insufficient saliva volume. Based on the duplicates, we calculated the mean cortisol concentration and log-transformed the values for all following analyses.

To obtain participants’ cortisol stress reactivity in response to the main part of the acute stress intervention, we calculated the difference between the cortisol value that was sampled directly prior to the acute stressor (at T_2_), which served as the pre-stress baseline, and the peak cortisol value post-stress (either at T_3_ or T_4_). We used the cortisol value at T_2_ as the pre-stress baseline because the initial sample (T_1_) is likely to be influenced by anticipatory HPA-axis activation, particularly in the case of MRI environments (Gossett et al., 2018; Noack et al., 2019). The T_2_ sample was collected immediately before the acute stress induction while participants had already acclimatized to the MRI environment, as has been done previously (Forbes et al., 2024). To account for interindividual variability in the time interval between sampling time points (because the exact duration of the object-location task runs was dependent on participants’ performance, leading to slight timing differences between individuals), the resulting differences were normalized to the time interval between sample collections. This peak-minus-baseline approach is consistent with established procedures in stress research (e.g., Miller et al., 2013; Schlotz et al., 2008).

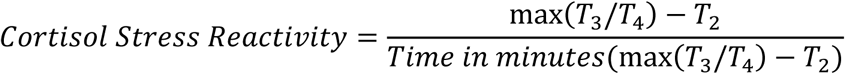

To determine how quickly participants’ post-stress cortisol decreased to their pre-stress cortisol levels (i.e., their stress recovery parameter), we calculated the difference between the abovementioned peak value (either at T_3_ or T_4_) and the last sampling timepoint following the acute stress phase on day 2 (T_5_), again, normalized to the time interval between sample collections:

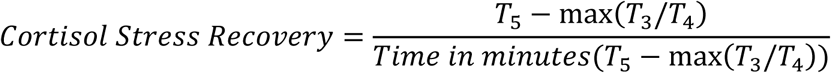

### 2.5. Subjective stress analysis

To quantify participants’ subjective stress reactivity in response to the acute stressor, we calculated the difference between the subjective stress rating obtained immediately before the acute stress phase (T_2_) and the peak subjective stress rating obtained after the stressor (either at T_3_ or T_4_). As above, to account for interindividual differences in the time interval between sampling time points, the difference was normalized by the time interval between sample collections:

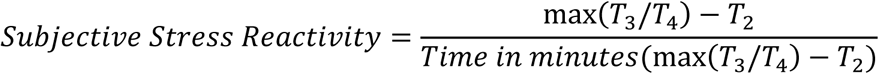

### 2.6. Stool sampling and analysis

Stool samples were self-sampled by participants using a home-collection kit (Sarstedt, Germany) that was provided to them on day 1 (including gloves, the sampling tube with a spatula, a gel cooling pad, and a Styrofoam container for transport). Participants were asked to freeze the sample and transport it back to the laboratory on study day 2, using the pre-cooled gel pad and Styrofoam container to prevent thawing. All participants were able to provide a stool sample during the study, and no participants were excluded or replaced due to difficulties with sample collection. Eighteen participants produced their stool samples on the evening of day 1, and the remaining 56 on day 2, before the experimental session that included the acute stress intervention (39 before 12:00 p.m. and 17 after 12:00 p.m.). Sampling times were similarly distributed across the stress and control groups with respect to both study day and time of day (see OSF: 10.17605/OSF.IO/RAVUY).

Samples were frozen (-25 °C) and, after all data collection was completed, were shipped to the Joint Microbiome Facility of the University of Vienna and the Medical University of Vienna, Austria (see details below). Along with sample collection, participants completed several questionnaires regarding their stool quality (Bristol Stool Chart; Lewis & and Heaton, 1997), dietary habits, and other lifestyle factors that can influence the gut microbiota (Johnson et al., 2020; Quigley, 2017), including screening questions that were partly based on the Human Microbiome Project (Core Microbiome Sampling Protocol A Version 4; https://commonfund.nih.gov/hmp). Participants of the stress and control groups showed comparable dietary patterns in terms of mainly animal-and plant-based diet, and only few individuals primarily consumed highly processed foods (*N*_total_ = 73, one participant was excluded due to missing dietary information, *n*_stress_ = 34; *n*_control_ = 39, Chi-square test: χ²(2) = 0.41, *p* = 0.81; Fisher’s exact *p* = 0.92; see also the OSF for a complete list of dietary-and lifestyle-related questions and the average values of stress/controls groups, 10.17605/OSF.IO/RAVUY).

In brief, DNA was extracted from 180-220 mg of stool using the QIAamp Fast DNA Stool Mini Kit on a QiaCube (Qiagen, The Netherlands) according to the manufacturer’s instructions. For microbial community profiling, the bacterial and archaeal V4 region of the 16S rRNA gene was amplified by polymerase chain reaction (PCR) and sequenced using primers 515F/806R (Apprill et al., 2015; Parada et al., 2016). DNA extraction, sequencing, and raw data processing were performed at the Joint Microbiome Facility (project ID: JMF-2212-18) as described previously (Pjevac et al., 2021), and sequencing was performed on an Illumina MiSeq (2 × 300 bp). Negative controls were included to monitor potential contamination. Amplicon pools were extracted from the raw sequencing data using the FASTQ workflow in BaseSpace (Illumina) with default parameters. Demultiplexing was performed with the Python package “demultiplex” (https://github.com/jfjlaros/demultiplex), allowing one mismatch for barcodes and two mismatches for linkers and primers. Amplicon sequence variants (ASVs) were inferred using the R-package “DADA2” (version 1.42; Callahan et al., 2016a), applying the recommended workflow ((Callahan et al., 2016b). FASTQ reads 1 and 2 were trimmed at 220/150 nt, with expected errors of 2. ASV sequences were subsequently classified using the classifier implemented in DADA2 against the SILVA database SSU Ref NR 99 (release 138.1; McLaren & Callahan, 2021; Quast et al., 2013) using a confidence threshold of 0.5. We used the SILVA database because it is actively maintained, extensively validated, and performs reliably across environments (McLaren & Callahan, 2021). If necessary, contaminants were removed *in silico* using the “decontam” software package (Davis et al., 2018). The raw 16S rRNA gene amplicon sequencing data is available at the Sequence Read Archive (https://www.ncbi.nlm.nih.gov/sra) under the BioProject accession PRJNA1366128.

To account for differences in sequencing depth and ensure comparability between individuals, we inspected the distribution of sequencing depths across all samples and rarefied the data to the minimum library size (5991). ASVs with a relative abundance < 0.1% were excluded from the samples to minimize the contribution of extremely rare sequences that may reflect sequencing noise. Because 16S rRNA gene amplicon sequencing data are compositional in nature (Bastiaanssen et al., 2023; Willis & Clausen, 2025), all gut microbial alpha diversity metrics were calculated from the rarefied ASV table. Alpha diversity indices (Shannon Index, Inverse Simpson Index, observed ASVs) rely on within-sample proportions and are therefore less sensitive to compositional artefacts than between-sample distance metrics (Gloor et al., 2017; Morton et al., 2017). Our preprocessing and analytical steps follow current recommendations for handling compositional 16S rRNA gene amplicon data (Knight et al., 2018; Sorbie et al., 2022; Willis & Clausen, 2025). To determine participants’ gut microbial community composition, ASV abundance data were analysed in terms of different alpha diversity metrics, such as Shannon Index, Inverse Simpson Index, and the number of observed ASVs (i.e., richness).

To quantify each individual’s inferred capacity to produce SCFAs, we used the same rarefied and filtered dataset as for all gut microbiota analyses (5991 reads per sample; low-abundance ASVs removed). SCFA-producing bacterial taxa were identified based on taxonomy-to-pathway mapping of SCFA-producing bacteria provided by Frolova et al. (2022). Their classification was derived from a metagenomic reconstruction of SCFA biosynthesis pathways across 2,856 reference genomes (see supplement, **Table S8**). Relative abundances were aggregated at the genus level, and genera were annotated as butyrate-or propionate-producers; acetate was not included because its biosynthesis pathway is highly abundant and present in almost all gut bacteria (Frolova et al., 2022; Louis & Flint, 2017). Because rare taxa were removed during preprocessing, SCFA estimates were based on genera with sufficient abundance to yield stable and biologically meaningful estimates (Weiss et al., 2017; Willis & Clausen, 2025). Some bacterial taxa possess multiple SCFA biosynthesis pathways for the same SCFA: these taxa were counted once, treating SCFA production as a binary phenotype at the genus level, consistent with the resolution of 16S rRNA gene amplicon data rather than a quantitative measure of pathway abundance. Inferred SCFA production capacity was then calculated as the summed relative abundance of all genera annotated as butyrate-or propionate-producers, yielding one aggregate value per SCFA per participant.

### 2.7. Statistical analysis

All analyses were conducted in R (version 4.4.0 R Core Team, 2024) in combination with “tidyverse” (version 2.0.0; Wickham et al., 2019) and Bioconductor (version 3.19; Huber et al., 2015). Microbiome data analysis was conducted using the “ampvis2” (version 2.8.3; Andersen et al., 2018) and “phyloseq” (version 1.48.0; McMurdie & Holmes, 2013) packages. Statistical tests and multiple comparison corrections were performed using “rstatix” (version 0.7.2; Alboukadel Kassambara, 2021). Separate independent-samples *t*-tests and analyses of variance (ANOVAs) were used to compare the groups and to analyze group × time interactions of salivary cortisol and subjective stress levels, respectively. Effect sizes for *t*-tests were calculated as Cohen’s *d* (Lakens, 2013). Unless stated otherwise, the α-level was set to 0.05 (two-tailed) for all analyses. Analyses were corrected for multiple comparisons when appropriate, using the Bonferroni-Holm correction. Results were visualized using “ggplot2” (version 3.5.1; Hadley, 2016), and model effects were illustrated using the “effects” package (version 4.2-2; Fox & Weisberg, 2018, 2019). Statistical tables were generated using the “stargazer” package (version 5.2.3; Hlavac, 2022). Raw data, analysis code, and the complete model outputs are openly available via the Open Science Framework (10.17605/OSF.IO/RAVUY).

#### 2.7.1. Model definition, evaluation, and comparison

To determine the effect of relevant predictors and covariates on cortisol stress reactivity, post-stress recovery, and subjective stress levels, we initially defined linear models (LMs) using the “stats” package (version 4.4.0; R Core Team, 2024b). However, due to violations of normality, heteroscedasticity, and to alleviate the effect of potential outliers, we subsequently used robust linear models (RLMs) with Huber’s loss function using the “MASS” package (version 7.3-61; Venables & Ripley, 2002). We computed a corresponding LM for each RLM to validate model stability and compared the explained variance (variance-based *R*² measure for RLMs), residual distributions, and sensitivity to outliers between models.

Model coefficients were tested for significance with the “coeftest” function from the “lmtest” package (version 0.9-40; Zeileis & Hothorn, 2002). To evaluate model fit, we calculated a variance-based *R*² measure for RLMs (using the “rlm” function from the “MASS” package), analogous to the coefficient of determination (*R*²) for ordinary least squares regression. The variance-based *R*² measure quantifies the proportion of variance explained by the model while down-weighting and accounting for the influence of potential outliers, computed as variance-based *R*² = 1-SS_res_ / SS_tot_ (where SS_res_ denotes the sum of squared residuals, and SS_tot_ denotes the total sum of squares, based on the mean value of the dependent variable). Additionally, we computed an adjusted variance-based *R*², which corrects for model complexity and sample size, providing a more conservative estimate of explained variance (Miles, 2005). The adjusted value was calculated as adjusted variance-based *R*^2^ = 1-(SS_res_ / SS_tot_) * ((*n*-1) / (*n*-*p*-1)), where *n* denotes the number of observations and *p* the number of predictors (excluding the intercept). RLM and LM comparisons were based on the unadjusted (variance-based) *R*² values, while adjusted variance-based *R*² values are reported for all results throughout the manuscript.

To ensure model robustness, we conducted outlier diagnostics using Cook’s distance (with the standard threshold of 4/n) and inspected the residuals. While a few observations exceeded this threshold, the robust estimation procedure inherently downweighted their influence (Huber, 1973; Verardi & Croux, 2009), ensuring that these values did not disproportionately affect the model coefficients. Robust linear models (RLMs) are known to support the stability and reliability of coefficient estimates in the presence of potential outliers (Wilcox, 2017). We then compared the RLMs to their corresponding LMs in terms of explained variance (variance-based *R*² measure for RLMs), residual distributions, and sensitivity to outliers. Across all models, RLMs explained more variance, residuals were more symmetrically distributed, Cook’s distances were markedly reduced, and skewness was lower compared to standard LMs. Note that model selection was based on data from the stress group (which was the group-of-interest), but that final gut microbiota models were also computed by incorporating both groups (to test for relevant group interactions) and by focusing on the control group only (to validate that potential effects were specific to the stress group but could not be found in controls).

#### 2.7.2. Selection of maximal and base models

Model selection followed a stepwise approach inspired by established guidelines for confirmatory modeling (Barr et al., 2013), beginning with a maximal model including all theoretically relevant predictors.

All covariates in the maximal model (biological sex, trait anxiety, perceived stress, and, where applicable, baseline cortisol, T_2_) were selected *a priori* based on established links to HPA-axis regulation and stress reactivity (Dickerson & Kemeny, 2004; Fiksdal et al., 2019; Kirschbaum & Hellhammer, 1994; Kudielka et al., 2009). Participants with known confounding factors for cortisol physiology or gut microbiota composition (e.g., recent antibiotic/probiotic use, gastrointestinal disorders, psychiatric or neurological diagnoses, hormonal contraceptive use) were already excluded during screening. Our microbiota predictors (gut microbial alpha diversity metrics, inferred SCFA production capacity) and gut microbiota-related variables were also defined *a priori* based on previous work (Cryan et al., 2019; Dalile et al., 2019, 2020; Foster et al., 2017; Kelly et al., 2016; Van De Wouw et al., 2018). This ensured that model evaluation was theory-driven and that microbial predictors were evaluated while controlling for all relevant psychophysiological covariates.

Although these recommendations were originally developed for mixed-effects models, we applied the same principled approach to robust linear models, simplifying them iteratively to achieve convergence and interpretability. Specifically, we began with a maximal model that predicted the three dependent variables (cortisol stress reactivity, post-stress recovery, subjective stress reactivity) using all relevant variables, including: biological sex, perceived stress (PSS; Cohen, 1988), trait anxiety (GAD-7; Spitzer et al., 2006), and baseline cortisol (T_2_) for cortisol stress reactivity and post-stress recovery.

### Maximal model

Dependent variable ∼ biological sex + trait anxiety + perceived stress + subjective stress + baseline cortisol (T_2_; the latter was only included for cortisol stress reactivity and post-stress recovery)

From the maximal model, non-significant variables were removed iteratively until the model converged and the maximum explained variance was achieved. This resulted in the base model that predicted the three dependent variables (cortisol stress reactivity, post-stress recovery, subjective stress reactivity).

### Base model

Dependent variable ∼ biological sex + trait anxiety + perceived stress + baseline cortisol (T_2_) (the latter was only included for cortisol stress reactivity and post-stress recovery)

Results pertaining to the base model revealed a (non-significant) effect of biological sex on cortisol stress reactivity, with biological males showing generally higher cortisol stress reactivity than biological females (*b* = 0.008, *SE* = 0.004, *z* = 1.92, *p* = 0.055). The model explained approximately 13.5% of the variance in cortisol reactivity (adjusted variance-based *R*² = 0.135). Turning to the post-stress recovery of salivary cortisol, biological males exhibited steeper cortisol recovery slopes than biological females (*b* =-0.002, *SE* = 0.002, *z* =-1.41, *p* = 0.158). Moreover, post-stress recovery of salivary cortisol was numerically (but not significantly) faster at higher levels of trait anxiety (*b* =-0.002, *SE* = 0.001, z =-1.86, *p* = 0.063) and significantly faster at higher baseline cortisol (*b* =-0.003, *SE* = 0.001, *z* =-3.43, *p* < 0.001). The predictors explained approximately 19.1% of the variance in post-stress recovery slopes (adjusted variance-based *R*² = 0.191). Finally, we examined predictors of subjective stress reactivity. In the base model, trait anxiety emerged as a significant predictor, with higher trait anxiety being associated with lower subjective stress responses (b =-0.044, SE = 0.022, z =-2.03, *p* = 0.042). Neither biological sex (*p* = 0.172) nor perceived stress (*p* = 0.154) was a significant predictor of subjective stress reactivity. The model explained approximately 5.6% of the variance in subjective stress reactivity (variance-based R² = 0.056; adjusted variance-based R² =-0.036).

Together, these base models allowed us to establish a reference point for evaluating the added explanatory value of gut microbiota parameters and microbiota-related variables in the following steps.

#### 2.7.3. Selection of gut microbiota models

Lastly, we defined separate RLMs that incorporated the gut microbial predictors and additional microbiota-related variables into the final base model. Similar to the approach described in section 2.7.2, we once again started from a maximal model that included gut microbiota predictors. Non-significant variables were iteratively removed until the model converged, and the maximum explained variance was achieved. This approach allowed us to examine whether variance in the dependent variables covaried with individual differences in gut microbiota composition across individuals. The gut microbial predictors included alpha diversity (either Shannon Index, Inverse Simpson Index, or the observed number of ASVs) and the microbiota-related variables such as diet, body-mass index (BMI), birth mode (vaginal or caesarean section) and early nutrition (breastfed, bottle-fed, or both). Finally, we used the same approach to assess how the relative abundance of SCFA-producing taxa covaried with the variance in the dependent variables across individuals.

### Gut microbiota model

Dependent variable ∼ gut microbiota measure (e.g., alpha diversity, SCFA-producing taxa) + biological sex + trait anxiety + perceived stress + baseline cortisol (T_2_) (the latter was only included for cortisol stress reactivity and post-stress recovery)

We also defined separate RLMs that included the Firmicutes/Bacteroidetes (F/B) ratio, as this has been linked to stress-related physiology and gut microbiota composition in previous studies (Jiang et al., 2015; Lach et al., 2018; Pusceddu et al., 2015). However, the F/B ratio did not emerge as a significant predictor of any of the stress-related outcome variables, and complete model outputs are available via the OSF (10.17605/OSF.IO/RAVUY).

## 3. Results

### 3.1. Acute stress intervention effectively increases salivary cortisol and subjective stress

Starting out, we verified that the stress and control groups were comparable in terms of age, biological sex, BMI (mean ± standard error of the mean, s.e.m.; stress group: 21.13 ± 0.40, controls: 22.08 ± 0.53), perceived stress (PSS; stress group: 18.17 ± 1.17, controls: 16.13 ± 1.11), and trait anxiety (GAD-7; stress group: 6.20 ± 0.67, controls: 5.72 ± 0.71; all *p* > 0.05). To test whether the acute stress induction affected stress-related measures, we conducted a repeated-measures ANOVA on the log-transformed salivary cortisol values with group (stress vs. control) as a between-subjects factor, and timepoint (7 sampling timepoints across both days) as a within-subjects factor. The analysis revealed a significant group × timepoint interaction (*F*(3.24, 233.53) = 3.647, *p* = 0.011, *η*² = 0.011; main effect of group: *F*(1.00, 72.00) = 6.068, *p* = 0.016, *η*² = 0.061; main effect of timepoint: *F*(3.24, 233.53) = 12.244, *p* < 0.001, *η*² = 0.037). As expected, post-hoc analysis showed comparable cortisol levels between the groups before acute stress induction (T_1_-T_2_; all *p* > 0.05), and significantly higher cortisol levels after stress in the stress group compared to controls (T_3_-T_6_; all *p* < 0.05, corrected for multiple comparisons, **Figure 1B**, **Table S1**, and see **Supplemental Figure S1** for an overview of the participant-specific cortisol trajectories).

Similarly, subjective stress levels showed a significant group × timepoint interaction (*F*(6, 420) = 2.440, *p* = 0.025, *η*² = 0.025), a significant main effect of timepoint (*F*(6, 420) = 3.025, *p* = 0.007, η² = 0.031), but no significant main effect of group (*F*(1, 70) = 1.661, *p* = 0.202, η² = 0.006). Post-hoc *t*-tests revealed comparable stress levels between the groups before acute stress induction (T_1_–T_2_; all *p* > 0.05) and significantly increased subjective stress in the stress group compared to the controls after stress induction (T3, *p* = 0.012, corrected for multiple comparisons), which recovered relatively quickly (T_4_-T_6_; all *p* > 0.005, **Figure 1C**, **Table S1**, and see **Supplemental Figure S1** for an overview of the participant-specific subjective stress levels). Together, these results indicate that the acute stress intervention successfully increased salivary cortisol and subjective stress levels in the stress group compared to the control group.

### 3.2. Gut microbial alpha diversity is associated with cortisol stress reactivity

Our main goal was to determine whether inter-individual differences in gut microbiota composition contributed to variations in participants’ responses to acute stress (Cryan et al., 2019; Foster et al., 2017). Specifically, we hypothesized that participants with higher gut microbial diversity would exhibit lower cortisol stress reactivity and a faster post-stress recovery. We thus tested whether three different metrics of gut microbial alpha diversity (Shannon Index, Inverse Simpson Index, observed number of ASVs) were associated with cortisol stress reactivity (alpha diversity appeared comparable between groups, separate independent-samples *t*-test, *n*_stress_ = 35 vs. *n*_control_ = 39, all *p* > 0.05). Cortisol stress reactivity was defined as the slope of cortisol increase following T_2_ (which noted the start of the acute stress intervention in the stress group; see Methods section for details). Our focus was on the stress group, but we also computed models that incorporated both groups (to test for potential group interactions) and that focused on the control group only (to verify that our expected effects were specific to the stress group but not present in controls).

Results showed a significant main effect of group in all three alpha diversity models (separate RLMs, *N*_total_ = 74, *n*_stress_ = 35, *n*_control_ = 39, all *p* < 0.001, see **Table 1** for complete model results), confirming higher cortisol stress reactivity in the stress group compared to controls. This was accompanied by significant alpha diversity × group interactions in all three models, indicating that the associations between gut microbial alpha diversity and cortisol stress reactivity were stronger and more positive in the stress group compared to controls (Shannon Index: *b* = 0.003, *SE* = 0.002, *z* = 2.07, *p* = 0.038, adjusted variance-based *R*² = 0.283; Inverse Simpson Index: *b* = 0.004, *SE* = 0.001, *z* = 2.65, *p* = 0.008, adjusted variance-based *R*² = 0.278; observed number of ASVs: *b* = 0.003, *SE* = 0.002, *z* = 2.17, *p* = 0.030, adjusted variance-based *R*² = 0.289; **Figure 2A**). In other words, higher gut microbial alpha diversity was more strongly associated with higher cortisol stress reactivity across those participants who had undergone the acute stress intervention. Biological sex showed a significant main effect in all three models (all *p* < 0.05), with biological males exhibiting generally higher cortisol stress reactivity than biological females. This difference was more pronounced in the stress group compared to controls (separate post-hoc RLMs to test for significant biological sex × group interactions, all *p* < 0.010; **Table S2**). Higher trait anxiety (all *p* < 0.001) was associated with higher cortisol stress reactivity in all models, along with higher levels of perceived stress that predicted lower cortisol reactivity across individuals (all *p* < 0.05, **Table 1**).

**Figure 2.**
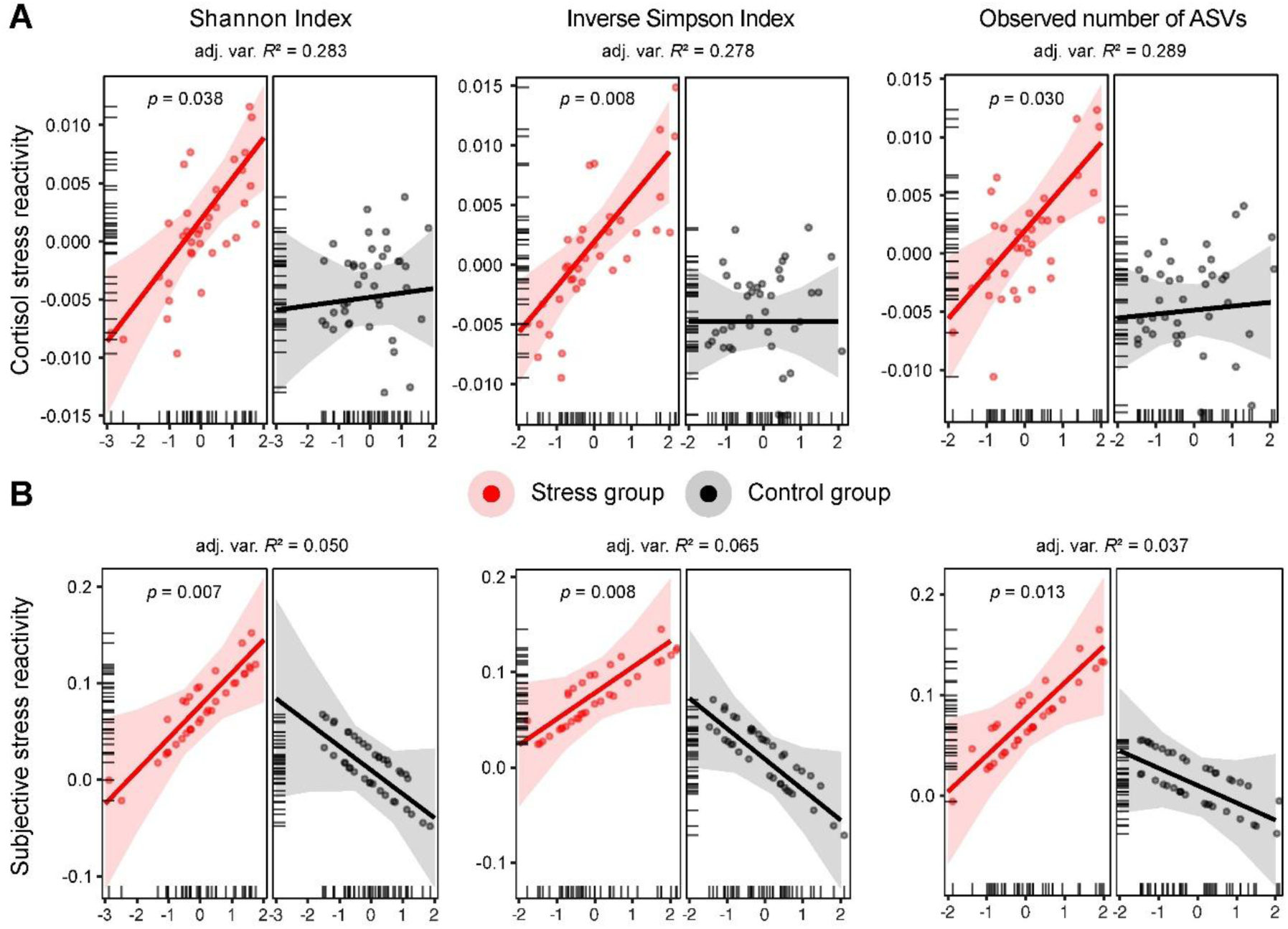
Gut microbial alpha diversity is associated with cortisol and subjective stress reactivity across groups. Partial effects from robust linear models (RLMs) predicting **(A)** cortisol stress reactivity (log-transformed concentrations, nmol/L) and **(B)** subjective stress reactivity based on gut microbial alpha diversity (Shannon Index, Inverse Simpson Index, observed number of ASVs) across the stress (red) and control (black) groups. Data points represent participant-specific fitted values; tick marks oriented toward the inside of the plot area indicate the distribution of the observed data points along the predictor axes; shaded areas around the regression line represent the 95% confidence intervals; *p*-values indicate significant gut microbial alpha diversity × group interaction effects; adjusted variance-based *R*2 values (shown as adj. var. *R*2) indicate overall model fit.

**Table 1.** Gut microbial alpha diversity predicting cortisol stress reactivity across groups. Results from robust linear regression models (RLMs) showed that cortisol stress reactivity across groups was predicted by gut microbial alpha diversity (Shannon Index, Inverse Simpson Index, observed number of ASVs), baseline cortisol, biological sex, and psychological variables (trait anxiety, perceived stress). Significance levels (Sig.): * *p* <.05, ** *p* <.005, *** *p* <.0001. Model formulas: Cortisol stress reactivity ∼ biological sex + trait anxiety + perceived stress + alpha diversity metric * group + baseline cortisol (T_2_).

| Predictor | *b* | SE | *z* | *p* | Sig. |
| --- | --- | --- | --- | --- | --- |
| *RLM (Shannon Index) predicting cortisol stress reactivity* |  |  |  |  |  |
| (Intercept) | -0.006 | 0.001 | -4.778 | 0 | *** |
| Biological sex | 0.003 | 0.002 | 2.215 | 0.027 | * |
| Trait anxiety | 0.004 | 0.001 | 3.511 | 0.0004 | *** |
| Perceived stress | -0.003 | 0.001 | -2.609 | 0.009 | ** |
| Shannon | 0.0004 | 0.001 | 0.315 | 0.753 |  |
| Stress group | 0.007 | 0.002 | 4.302 | 0.00002 | *** |
| Baseline cortisol (T<sub>2</sub>) | -0.002 | 0.001 | -2.4 | 0.016 | * |
| Shannon:Stress group | 0.003 | 0.002 | 2.07 | 0.038 | * |
| *RLM (Inverse Simpson Index) predicting cortisol stress reactivity* |  |  |  |  |  |
| (Intercept) | -0.006 | 0.001 | -4.973 | 0 | *** |
| Biological sex | 0.003 | 0.001 | 2.113 | 0.035 | * |
| Trait anxiety | 0.005 | 0.001 | 3.937 | 0.0001 | *** |
| Perceived stress | -0.004 | 0.001 | -2.964 | 0.003 | ** |
| Inverse Simpson | 0 | 0.001 | -0.001 | 1 |  |
| Stress group | 0.007 | 0.001 | 4.601 | 0 | *** |
| Baseline cortisol (T<sub>2</sub>) | -0.002 | 0.001 | -2.223 | 0.026 | * |
| Inverse Simpson:Stress group | 0.004 | 0.001 | 2.654 | 0.008 | ** |
| *RLM (observed number of ASVs) predicting cortisol stress reactivity* |  |  |  |  |  |
| (Intercept) | -0.006 | 0.001 | -4.617 | 0 | *** |
| Biological sex | 0.003 | 0.002 | 2.128 | 0.033 | * |
| Trait anxiety | 0.005 | 0.001 | 3.478 | 0.001 | *** |
| Perceived stress | -0.003 | 0.001 | -2.538 | 0.011 | * |
| Observed | 0.0003 | 0.001 | 0.318 | 0.751 |  |
| Stress group | 0.007 | 0.002 | 4.175 | 0.00003 | *** |
| Baseline cortisol (T<sub>2</sub>) | -0.002 | 0.001 | -2.515 | 0.012 | * |
| Observed:Stress group | 0.003 | 0.002 | 2.168 | 0.03 | * |

To break down these complex models, we repeated the analyses for the stress group only (separate RLMs, *n_stress_* = 35). Higher gut microbial alpha diversity was tied to higher cortisol stress reactivity across participants (Shannon Index: *b* = 0.004, *SE* = 0.001, *z* = 3.15, *p* = 0.002; adjusted variance-based *R*² = 0.243; Inverse Simpson Index: *b* = 0.004, *SE* = 0.001, *z* = 3.45, *p* = 0.001; adjusted variance-based *R*² = 0.249; observed number of ASVs: *b* = 0.004, *SE* = 0.002, *z* = 2.58, *p* = 0.01; adjusted variance-based *R*² = 0.253; **Figure S2**, and see **Table S3** for complete model results; see also **Supplementary Results 1** showing that stool quality did not explain the effects). Again, biological males showed generally higher cortisol stress reactivity in all three alpha diversity models (all *p* < 0.05; **Figure S2**; and see **Supplementary Results 2-3** for additional analyses). Higher trait anxiety was associated with higher cortisol stress reactivity in the Inverse Simpson and observed number of ASVs models (*p* < 0.05). Importantly, we did not detect any significant relationships between any of the three alpha diversity metrics and cortisol stress reactivity in the control group (separate RLMs, *n*_control_ = 39, all *p* > 0.05), verifying that the observed effects were specific to the acute stress intervention (complete model outputs are available via the OSF, 10.17605/OSF.IO/RAVUY).

Next, we reasoned that gut microbiota diversity may not only buffer against inadequate (i.e., blunted or exaggerated) stress reactivity but that it also may support faster post-stress recovery, which reflects how efficiently HPA axis activity returns to homeostasis in the aftermath of stress (McEwen, 1993; Sapolsky et al., 2000; Sudo et al., 2004). To capture this, we quantified post-stress recovery as the slope of cortisol decline after the stress peak (see Methods section for details). We then repeated the abovementioned analysis and examined the relationship between the three different alpha diversity metrics and post-stress recovery across groups (separate RLMs, *N*_total_ = 74) and separately for each group (separate RLMs, *n*_stress_ = 35, *n*_control_ = 39). In contrast to cortisol stress reactivity, none of the alpha diversity metrics were significantly associated with post-stress recovery in any of the models (all *p* > 0.05, **Tables S4-5**).

To compare our findings with alternative measures that describe cortisol dynamics, we also modelled cortisol responses as the area under the curve (AUC) with respect to increase (AUCi) and ground (AUCg; collapsing across cortisol stress reactivity and post-stress recovery timepoints; Pruessner et al., 2003). This essentially replicated the main findings, showing that higher gut microbial alpha diversity was linked to higher AUCi and AUCg in the stress group (**Supplementary Results 4**). We also examined cortisol dynamics on a per-timepoint basis, which confirmed the results once more (**Supplementary Results 5**).

Taken together, and contrary to our expectations, results indicated that higher gut microbial alpha diversity was significantly associated with higher cortisol stress reactivity in the stress group but not in controls. No significant associations were observed between gut microbial alpha diversity and post-stress recovery.

### 3.3. Gut microbial alpha diversity is associated with subjective stress reactivity

Because cortisol dynamics in response to acute stress do not necessarily reflect subjectively experienced stress (Campbell & Ehlert, 2012), we next evaluated whether the relationship between gut microbial alpha diversity and stress reactivity was also mirrored in terms of subjective stress levels. To this end, we quantified subjective stress reactivity similarly to cortisol stress reactivity (i.e., the slope of the increase in subjective stress levels following T_2_, which noted the start of the acute stress intervention in the stress group) and repeated the analysis using subjective stress reactivity as the dependent variable (see Methods section for details).

Results revealed a significant main effect of group in all three alpha diversity models (separate RLMs, *N*_total_ = 74, *n*_stress_ = 35; *n*_control_ = 39, all *p* < 0.01, see **Table 2** for complete model results), confirming higher subjective stress reactivity in the stress group compared to the controls. This was accompanied by significant alpha diversity × group interactions in all three models (all *p* < 0.05), indicating that the associations between gut microbial alpha diversity and subjective stress reactivity were stronger and more positive in the stress group compared to the control group (Shannon Index: *b* = 0.059, *SE* = 0.022, *z* = 2.68, *p* = 0.007; adjusted variance-based *R*² = 0.050; Inverse Simpson Index: *b* = 0.059, *SE* = 0.022, *z* = 2.67, *p* = 0.008; adjusted variance-based *R*² = 0.065; observed number of ASVs: *b* = 0.053, *SE* = 0.022, *z* = 2.47, *p* = 0.013; adjusted variance-based *R*² = 0.037; **Figure 2B**). In other words, higher gut microbial alpha diversity was more strongly associated with higher subjective stress reactivity across those participants who had undergone the acute stress intervention.

**Table 2.**
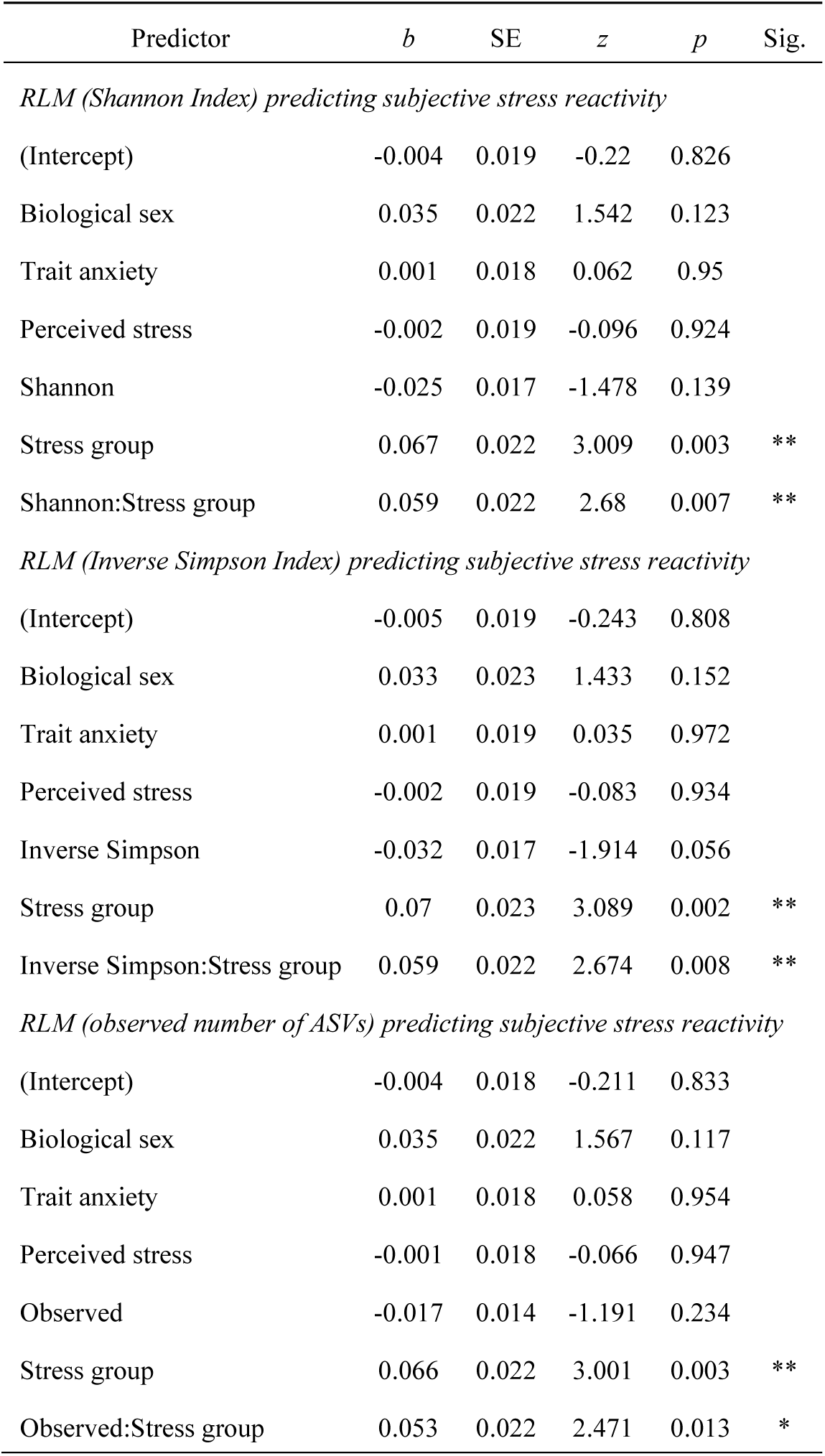
Gut microbial alpha diversity predicting subjective stress reactivity across groups. Results from robust linear regression models (RLMs) showed that subjective stress reactivity across groups was predicted by gut microbial alpha diversity (Shannon Index, Inverse Simpson Index, observed number of ASVs), biological sex, and psychological variables (trait anxiety, perceived stress). Significance levels (Sig.): * *p* <.05, ** *p* <.005, *** *p* <.0001. Model formulas: Subjective stress reactivity ∼ biological sex + trait anxiety + perceived stress + alpha diversity metric * group.

| Predictor | *b* | SE | *z* | *p* | Sig. |
| --- | --- | --- | --- | --- | --- |
| *RLM (Shannon Index) predicting subjective stress reactivity* |  |  |  |  |  |
| (Intercept) | -0.004 | 0.019 | -0.22 | 0.826 |  |
| Biological sex | 0.035 | 0.022 | 1.542 | 0.123 |  |
| Trait anxiety | 0.001 | 0.018 | 0.062 | 0.95 |  |
| Perceived stress | -0.002 | 0.019 | -0.096 | 0.924 |  |
| Shannon | -0.025 | 0.017 | -1.478 | 0.139 |  |
| Stress group | 0.067 | 0.022 | 3.009 | 0.003 | ** |
| Shannon:Stress group | 0.059 | 0.022 | 2.68 | 0.007 | ** |
| *RLM (Inverse Simpson Index) predicting subjective stress reactivity* |  |  |  |  |  |
| (Intercept) | -0.005 | 0.019 | -0.243 | 0.808 |  |
| Biological sex | 0.033 | 0.023 | 1.433 | 0.152 |  |
| Trait anxiety | 0.001 | 0.019 | 0.035 | 0.972 |  |
| Perceived stress | -0.002 | 0.019 | -0.083 | 0.934 |  |
| Inverse Simpson | -0.032 | 0.017 | -1.914 | 0.056 |  |
| Stress group | 0.07 | 0.023 | 3.089 | 0.002 | ** |
| Inverse Simpson:Stress group | 0.059 | 0.022 | 2.674 | 0.008 | ** |
| *RLM (observed number of ASVs) predicting subjective stress reactivity* |  |  |  |  |  |
| (Intercept) | -0.004 | 0.018 | -0.211 | 0.833 |  |
| Biological sex | 0.035 | 0.022 | 1.567 | 0.117 |  |
| Trait anxiety | 0.001 | 0.018 | 0.058 | 0.954 |  |
| Perceived stress | -0.001 | 0.018 | -0.066 | 0.947 |  |
| Observed | -0.017 | 0.014 | -1.191 | 0.234 |  |
| Stress group | 0.066 | 0.022 | 3.001 | 0.003 | ** |
| Observed:Stress group | 0.053 | 0.022 | 2.471 | 0.013 | * |

We then broke down these models and repeated the analyses for the stress group only (separate RLMs, *n_stress_* = 35). Higher gut microbial alpha diversity was significantly associated with higher subjective stress reactivity across participants (Shannon Index: *b* = 0.028, *SE* = 0.012, *z* = 2.42, *p* = 0.015; adjusted variance-based *R*² = 0.028; Inverse Simpson Index: *b* = 0.024, *SE* = 0.012, *z* = 2.07, *p* = 0.039; variance-based *R*² = 0.116; adjusted variance-based *R*² =-0.002; observed number of ASVs: *b* = 0.033, *SE* = 0.013, *z* = 2.49, *p* = 0.013; adjusted variance-based *R*² = 0.034; **Figure S2** and **Table S6**; see also **Supplementary Results 1** showing that stool quality did not explain the effects). Across models, male biological sex was significantly associated with higher subjective stress reactivity in the Shannon model (*p* = 0.045), but not in the Inverse Simpson (*p* = 0.072) or observed number of ASVs models (*p* = 0.054; **Figure S2)**. Trait anxiety and perceived stress was not significantly associated with subjective stress reactivity in any of the models (all *p* > 0.05; **Table S6**). We also conducted an additional analysis assessing subjective post-stress recovery (analogous to the analysis of post-stress recovery of cortisol trajectories above) but found no significant effects (all *p* > 0.05; **Table S7**). Importantly, we did not detect any significant relationships between the tested gut microbial alpha diversity metrics and subjective stress reactivity in the control group (separate RLMs, *n*_control_ = 39, all *p* > 0.05), verifying that the observed effects were specific to the acute stress intervention.

Thus, in line with our results on cortisol stress reactivity, we found that higher gut microbial alpha diversity was significantly associated with higher subjective stress reactivity in the stress group but not in the control group.

### 3.4. Inferred gut microbial capacity to produce butyrate and propionate is associated with cortisol stress reactivity, but not subjectively experienced stress

The gut microbiota was shown to influence stress physiology through the production of short-chain fatty acids (SCFAs). Prior evidence highlighted that SCFAs (particularly butyrate, but also propionate and acetate) reduced corticosterone release following acute stress exposure in chronically stressed mice (Van De Wouw et al., 2018), and that SCFAs attenuated the cortisol surge after psychological stress in humans (Dalile et al., 2020). We hypothesized that a higher microbial capacity to produce SCFAs would be tied to lower cortisol stress reactivity and faster post-stress recovery. We thus estimated each individual’s microbial SCFA production capacity by identifying bacterial taxa with known butyrate-or propionate-producing capabilities based on their taxonomic classifications (see Methods section for details; **Table S8**). We did not include acetate in our analysis, as its biosynthesis pathway is highly abundant and present in almost all gut bacteria (Frolova et al., 2022; Louis & Flint, 2017). We then grouped bacterial taxa according to their capacity to produce butyrate and propionate and calculated the summed relative abundance per participant (thus, yielding estimates of the relative abundance of potential butyrate-and propionate-producing bacteria in each participant’s gut microbiota).

Across the participants in the stress group, we identified butyrate-producing bacteria with a mean relative abundance of 29.43% (range: 12.25-54.90%). These values are consistent with estimates from Frolova et al. (2022), who predicted SCFA production phenotypes based on a large-scale twin-cohort (*N* = 3288; Goodrich et al., 2016), and who reported a mean predicted butyrate-producing phenotype abundance of 28.0% (IQR: 21.0-34.3%). In the control group, the mean relative abundance of butyrate-producing bacteria was 30.75% (range: 16.66-48.72%). Propionate-producing bacteria showed a mean relative abundance of 89.40% in the stress group (range: 73.83-97.16%) and 87.81% in controls (range: 70.68-97.70%), which is substantially higher than expected based on the previously reported mean of 35.9% (Frolova et al., 2022; but note that this discrepancy may reflect differences in the taxonomic classification). As expected, there were no significant differences in butyrate-or propionate-producing bacteria between the groups (separate independent-samples *t*-test, *n*_stress_ = 35 vs. *n*_control_ = 39, all *p* > 0.05). We next tested whether the relationship between the inferred gut microbial SCFA production capacity (butyrate, propionate) and the stress-related measures (cortisol/subjective stress reactivity/post-stress recovery) was specific to acute stress.

Results revealed a significant main effect of group in both SCFA models (separate RLMs, *N*_total_ = 74, *n*_stress_ = 35, *n*_control_ = 39, all *p* < 0.001; see **Table 3** for complete model results), confirming higher cortisol stress reactivity in the stress group compared to controls. Significant interaction effects indicated that the associations between inferred SCFA production capacity and cortisol stress reactivity differed between the groups (butyrate: *b* = 0.004, *SE* = 0.002, *z* = 2.11, *p* = 0.035, adjusted variance-based *R*² = 0.272; propionate: *b* =-0.003, *SE* = 0.002, *z* =-2.14, *p* = 0.033, adjusted variance-based *R*² = 0.249; **Figure 3**). In other words, a higher relative abundance of potential butyrate producers was more strongly tied to higher cortisol stress reactivity in the stress compared to the control group. The opposite was the case for propionate, where a higher relative abundance of potential propionate producers was more strongly tied to lower cortisol stress reactivity in the stress group compared to controls.

**Figure 3.**
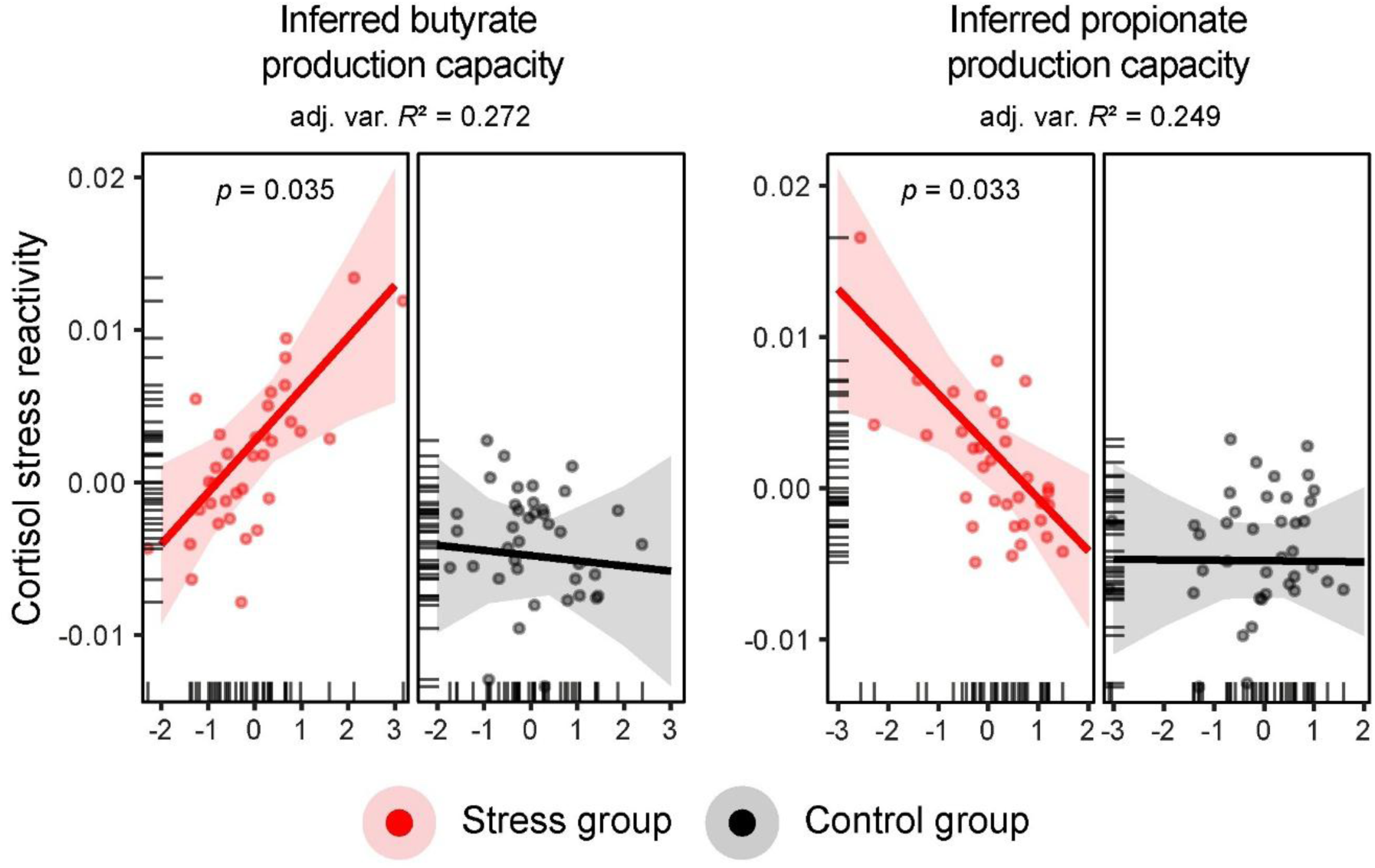
Inferred gut microbial capacity to produce butyrate and propionate is associated with cortisol stress reactivity across groups. Partial effects from robust linear models (RLMs) predicting cortisol stress reactivity (log-transformed concentrations, nmol/L) based on inferred SCFA production capacity for butyrate (left panel) and propionate (right panel) across the stress (red) and control (black) groups. Data points represent participant-specific fitted values; tick marks oriented toward the inside of the plot area indicate the distribution of the observed data points along the predictor axes; shaded areas around the regression line represent the 95% confidence intervals; *p*-values indicate significant inferred SCFA production capacity × group interaction effects; adjusted variance-based *R*2 values (shown as adj. var. *R*2) indicate overall model fit.

**Table 3.** Inferred gut microbial capacity to produce butyrate and propionate predicting cortisol stress reactivity across groups. Results from robust linear regression models (RLMs) predicting cortisol stress reactivity across groups based on the inferred gut microbial capacity to produce SCFAs (butyrate, propionate), baseline cortisol, biological sex, and psychological variables (trait anxiety, perceived stress). Significance levels (Sig.): * *p* <.05, ** *p* <.005, *** *p* <.0001. Model formulas: Cortisol stress reactivity ∼ biological sex + trait anxiety + perceived stress + SCFAs * group + baseline cortisol (T₂).

| Predictor | *b* | SE | *z* | *p* | Sig. |
| --- | --- | --- | --- | --- | --- |
| *RLM (butyrate) predicting cortisol stress reactivity* |  |  |  |  |  |
| (Intercept) | -0.006 | 0.002 | -4.062 | 0.00005 | *** |
| Biological sex | 0.003 | 0.002 | 1.799 | 0.072 |  |
| Trait anxiety | 0.004 | 0.001 | 2.436 | 0.015 | * |
| Perceived Stress | -0.002 | 0.001 | -1.461 | 0.144 |  |
| Butyrate | -0.0003 | 0.001 | -0.27 | 0.788 |  |
| Stress group | 0.008 | 0.002 | 4.199 | 0.00003 | *** |
| Baseline cortisol (T<sub>2</sub>) | -0.002 | 0.001 | -2.526 | 0.012 | * |
| Butyrate:Stress group | 0.004 | 0.002 | 2.106 | 0.035 | * |
| *RLM (propionate) predicting cortisol stress reactivity* |  |  |  |  |  |
| (Intercept) | -0.006 | 0.001 | -4.355 | 0.00001 | *** |
| Biological sex | 0.003 | 0.002 | 1.755 | 0.079 |  |
| Trait anxiety | 0.005 | 0.001 | 3.588 | 0.0003 | *** |
| Perceived Stress | -0.003 | 0.001 | -2.464 | 0.014 | * |
| Propionate | -0.00003 | 0.001 | -0.032 | 0.974 |  |
| Stress group | 0.008 | 0.002 | 4.583 | 0 | *** |
| Baseline cortisol (T<sub>2</sub>) | -0.002 | 0.001 | -2.225 | 0.026 | * |
| Propionate:Stress group | -0.003 | 0.002 | -2.136 | 0.033 | * |

When focusing on the stress group alone, a numerically higher (but non-significant) relative abundance of butyrate-producing bacteria appeared associated with higher cortisol stress reactivity (separate RLMs, *n*_stress_ = 35, *b* = 0.0035, *SE* = 0.0018, *z* = 1.91, *p* = 0.056; adjusted variance-based *R*² = 0.173; **Figure S3**, **Table S9**), while propionate-producing bacteria were significantly associated with lower cortisol stress reactivity across individuals (*b* =-0.0038, *SE* = 0.0018, *z* =-2.02, *p* = 0.042; adjusted variance-based *R*² = 0.176; **Figure S3**, **Table S9**; see also **Supplementary Results 1** showing that stool quality did not explain the effects). When including both SCFAs in the same model, butyrate (*b* = 0.0041, *SE* = 0.0014, *z* = 2.88, *p* = 0.004) and propionate (*b* =-0.0049, *SE* = 0.0017, *z* =-2.98, *p* = 0.003) remained significant predictors of cortisol stress reactivity (adjusted variance-based *R*² = 0.278; **Table S9**). There were no significant associations between the inferred SCFA production capacity and any other (cortisol or subjective) stress-related measures (all *p* > 0.05; **Tables S10-12**). Importantly, there were no significant relationships between inferred SCFA production capacity and any of the stress-related measures in controls (separate RLMs, *n*_control_ = 39, all *p* > 0.05). Once again, we validated the findings using AUC-related measures and by examining the SCFA × stress relationship on a per-timepoint basis, which gave virtually identical results (**Supplementary Results 4-5**).

To summarize, we found that a higher relative abundance of potential butyrate-producing bacteria was specifically tied to higher cortisol stress reactivity across participants, whereas a higher relative abundance of potential propionate-producing bacteria was associated with lower cortisol stress reactivity. These effects appeared specific to cortisol stress reactivity (but not subjective stress reactivity or post-stress recovery) in the stress group, and no effects were observed in controls.

## 4. Discussion

In the present study, we investigated whether individual differences in gut microbiota composition and in the inferred gut microbial capacity to produce certain SCFAs were linked to (cortisol and subjective) stress reactivity and post-stress recovery following acute stress in healthy adults. Several key findings emerged: First, higher gut microbial alpha diversity was associated with higher cortisol stress reactivity and higher levels of subjective stress across individuals. This effect was contrary to what we had expected but was consistently observed for several well-established alpha diversity metrics, highlighting a modulatory role of the gut microbiota on the physiological and psychological dimensions of stress. Surprisingly, we were unable to confirm a relationship between gut microbial alpha diversity and post-stress recovery. Second, individuals with a higher relative abundance of butyrate-producing bacteria tended to show higher cortisol stress reactivity in response to acute stress. This relationship was reversed for propionate, where a higher relative abundance of propionate-producing bacteria was tied to lower cortisol stress reactivity. Together, these results are the first to highlight the connection between gut microbiota composition, inferred SCFA production capacity, and the psychophysiological response to acute stress in healthy adults.

Contrary to what we had hypothesized, we found that higher gut microbial alpha diversity was associated with higher (instead of lower) cortisol stress reactivity across individuals who underwent an acute stress intervention. This association persisted when controlling for potential confounding factors (such as baseline cortisol or trait anxiety) and remained consistent across different alpha diversity metrics (Shannon Index, Inverse Simpson Index, observed number of ASVs). We interpret this relationship as gut microbial diversity being associated with adequate, time-limited cortisol stress reactivity, which reflects an important marker of a flexible and adaptive stress response (Kudielka et al., 2009; McEwen, 2017; Miller et al., 2007). Prior animal research focused on extreme imbalances in HPA-axis activity, reporting an exaggerated HPA axis response in stress-exposed germ-free mice. The exaggerated response was normalized with probiotic colonization that increased microbial diversity and SCFA production (Sudo et al., 2004). Similarly, probiotic administration reduced corticosterone release and anxiety-like behavior in animals (Bravo et al., 2011). Only a few studies have explored the gut microbiota-stress link in humans, mainly focusing on probiotic supplementation (for example, highlighting that probiotics may buffer against the negative effects of acute stress on cognition and mood; Papalini et al., 2019; Steenbergen et al., 2015), on patients with major depression (Kelly et al., 2016) or infants (Rosin et al., 2021). More recently, Boehme et al. (2023) reported probiotic effects on cortisol dynamics following an acute stressor; however, their analysis focused on overall cortisol output (area under the curve) rather than phasic cortisol stress reactivity or post-stress recovery, and did not examine gut microbiota composition beyond the supplemented strain. Schmidt et al. (2015) reported prebiotic effects on the cortisol awakening response, which is a circadian-driven marker of trait-like HPA-axis regulation different from the phasic cortisol stress reactivity examined here. Another important aspect is that cortisol dynamics and subjectively experienced stress are often uncorrelated (Campbell & Ehlert, 2012; Hellhammer & Schubert, 2012), with some individuals exhibiting elevated cortisol levels without reporting elevated subjective stress or vice versa. In line with this dissociation, subjective stress ratings peaked earlier (at T_3_) than salivary cortisol concentrations (at T_4_). This temporal offset is well-established, as subjective stress may reflect the immediate psychological appraisal of the stressor (Campbell & Ehlert, 2012; Hellhammer & Schubert, 2012), whereas cortisol secretion follows HPA-axis dynamics and typically peaks 20-30 minutes after stress onset (Dickerson & Kemeny, 2004; Linz et al., 2019). Here, we found that higher gut microbial alpha diversity was also associated with higher subjective stress reactivity. Put differently, individuals with a more diverse gut microbiota composition were highly responsive in their psychophysiological parameters when challenged by a standardized acute stress intervention. We also examined the role of biological sex since both the acute stress response and gut microbiota composition were shown to differ depending on biological sex (Adjei et al., 2018; Stefanaki et al., 2022). While we found that biological males exhibited generally higher cortisol stress reactivity than females, we observed a sex-dependent effect of gut microbial alpha diversity (Shannon Index) on subjective stress reactivity. Overall, we present the first evidence in healthy adults that underscores the potential buffering role of a diverse gut microbiota in the flexible response to acute stress.

From a mechanistic viewpoint, higher alpha diversity was proposed to reflect a more stable microbial ecosystem that confers functional resilience through taxonomic redundancy (Lozupone et al., 2012), potentially buffering against perturbations (Shade et al., 2012), such as psychological stress (Foster & Neufeld, 2013; Sheng et al., 2025). Altered microbiota diversity was reported in patients with major depression (Kelly et al., 2016; Zheng et al., 2016), anxiety disorders (Jiang et al., 2015, 2018), and irritable bowel disease (Pittayanon et al., 2019). Reduced alpha diversity can co-occur with elevated gut permeability, as demonstrated in alcohol-dependent individuals (Leclercq et al., 2014), facilitating the translocation of bacterial components such as lipopolysaccharides into the circulation (Madison & Bailey, 2024). This, in turn, can trigger systemic inflammation and chronic HPA axis activation (Kelly et al., 2015; Madison & Bailey, 2024), which is considered detrimental compared to the time-limited HPA axis activity in response to acute stress. By preserving intestinal barrier integrity, a diverse gut microbiota may help suppress such systemic inflammation, keeping neuroinflammation and the chronic activation of stress-related brain circuits in check (Madison & Bailey, 2024). For instance, gut microbiota depletion in mice was shown to disrupt the transcriptomic and metabolic profiles of stress-sensitive brain regions (such as the hippocampus and amygdala), resulting in exaggerated HPA axis activity and altered stress responsivity (Tofani, Leigh, et al., 2025). This establishes the gut microbiota as an important facilitator of the stress response. Here, we show for the first time that the gut microbiota is tied to (cortisol and subjective) stress reactivity also in healthy adults.

Higher microbial diversity may also enhance an organism’s metabolic flexibility (Moya & Ferrer, 2016), boosting the production of neuroactive metabolites (Bourassa et al., 2016; Dalile et al., 2019; Rea et al., 2016). SCFAs are produced by the microbial fermentation of dietary fibers, help to preserve intestinal barrier integrity, regulate immune function, and can affect the central nervous system via vagal and epigenetic pathways (Cryan et al., 2019; Dalile et al., 2019; Peng et al., 2009; Smith et al., 2013; Stilling et al., 2016). We focused our investigation on two of the most common SCFAs, butyrate and propionate (we excluded acetate due to its ubiquitous biosynthesis pathway; Frolova et al., 2022; Louis & Flint, 2017), and hypothesized that a higher inferred gut microbial capacity to produce these SCFAs would be associated with a diminished effect of acute stress across individuals. Different from what we had predicted (but in line with our abovementioned findings regarding cortisol and subjective stress reactivity), we found that a higher relative abundance of potential butyrate-producing bacteria was correlated with higher cortisol stress reactivity (but not subjective stress reactivity or post-stress recovery). At a first glance, this result may appear contrary to previous studies in rodents that demonstrated reduced stress-related corticosterone release after supplementation with a mixture of SCFAs (Van De Wouw et al., 2018) or with butyrate alone (Wang et al., 2023). However, a more detailed analysis of their findings reveals that SCFA supplementation lowered *overall* levels of corticosterone release (rather than specific effects related to acute stress reactivity, as we tested here), suggesting that SCFA supplementation could still have caused higher corticosterone reactivity to acute stress. In humans, SCFA supplementation was linked to a more nuanced effect on cortisol under psychosocial stress, with both doses attenuating cortisol levels compared to placebo (Dalile et al., 2020). However, this effect was not replicated in a follow-up study that only administered butyrate (Dalile et al., 2024). These inconsistencies likely reflect differences between exogenous SCFA supplementation and endogenous SCFA production (as was the focus here). Furthermore, an individual’s capacity to produce SCFAs appears directly tied to gut microbial diversity (Frolova et al., 2022), as more diverse communities can occupy a broader array of metabolic niches and entertain cross-feeding interactions (Moya & Ferrer, 2016). For example, the butyrate-producing bacteria *Faecalibacterium prausnitzii* and *Roseburia* spp. rely on acetate or lactate production from other microbes, illustrating metabolic interdependence within microbial communities. Thus, a diverse and functionally redundant microbial community can likely sustain SCFA production and exert beneficial effects on HPA axis activity.

Unlike butyrate, we found that the inferred capacity to produce propionate was associated with lower cortisol stress reactivity across individuals who had undergone the acute stress intervention. This is in line with prior studies that reported mixed results when comparing propionate and butyrate. While some studies demonstrated attenuated cortisol dynamics of propionate supplementation on stress physiology (Dalile et al., 2020; Van De Wouw et al., 2018), others noted no or even anxiogenic effects at higher propionate concentrations (Choi et al., 2018; Kimura et al., 2011). It is a known phenomenon that SCFAs can exert distinct and even opposing effects on host physiology (Kimura et al., 2011). Their stress-related actions may depend on the exact mixture and dosage of supplementation, the type of physiological system that is primarily modulated (e.g., immune-, endocrine-, or neural systems), or host specifics (e.g., interindividual differences in microbiota composition, or stress sensitivity; Dalile et al., 2020; Rosell-Cardona et al., 2025). Together, these findings position endogenous SCFA production capacity, particularly related to butyrate and propionate, as candidate pathways that link the gut microbiota with HPA axis regulation.

Specific pathways may include direct actions of SCFAs via receptor-binding to free fatty acid receptors (FFAR2/3) on enteroendocrine and immune cells (Maslowski et al., 2009; Tedelind et al., 2007), as well as to vagal afferents that relay microbial signals to stress-related brain regions (De Vadder et al., 2014; Fülling et al., 2019; Kimura et al., 2011). SCFAs can further cross the blood-brain barrier and inhibit histone deacetylases, affecting stress-related gene expression (Lupori et al., 2022; Stilling et al., 2016). Indirectly, SCFAs were shown to reduce inflammation, T-cells, and microglia (Silva et al., 2020; Smith et al., 2013; Wenzel et al., 2020). Butyrate in particular preserves barrier integrity by preventing bacterial translocation under stress (Braniste et al., 2014; Kelly et al., 2015; Rosell-Cardona et al., 2025). Here, we found that a higher inferred capacity to produce butyrate and propionate was associated with cortisol stress reactivity. While we are unable to pinpoint the specific molecular mechanisms, we suggest that gut microbial SCFA producers may influence the psychophysiological response to acute stress via multiple mechanisms, ranging from direct interactions with the HPA-axis to indirect effects on barrier integrity and neuroinflammation.

Recent work highlights that gut microbiota-stress associations involve not only compositional differences but also dynamic properties of the microbial ecosystem. Bastiaanssen et al. (2021) showed that greater microbiota volatility (thus, describing temporal instability, or the degree of compositional change across timepoints), predicts stronger hormonal and behavioral stress responses in both mice and humans, despite species-specific taxonomic patterns. They also reported functional convergence across organisms, with conserved metabolic pathways underlying stress-related phenotypes. This supports our interpretation that SCFA-related pathways may represent stable microbial features that are relevant to stress physiology, even if taxonomic profiles may diverge between species. Unfortunately, our study relied only on a single stool sample per participant, which is why were unable to quantify volatility (thus, results represent a compositional snapshot rather than a dynamic profile). More broadly, associations between microbiota composition, cortisol responses, and anxiety-like behavior are not necessarily consistent across model organisms, reflecting differences in host physiology, microbial ecology, and stress paradigms (Nagpal & Cryan, 2021). This cross-species variability suggests that gut microbiota-stress links are state-dependent and that not all associations may generalize across baseline and stress conditions, which is consistent with our observation that microbiota-stress associations were detectable in the stress group but not in controls.

Another important aspect is that circadian rhythms are known to influence both cortisol secretion and gut microbiota composition (Kiessling et al., 2025; Liu, 2024; Tofani, Clarke, et al., 2025). Here, experimental sessions were conducted in the afternoons to minimize diurnal variability in cortisol trajectories (Dickerson & Kemeny, 2004). Although circadian oscillations of microbial taxa and SCFA production are well-documented (Kiessling et al., 2025; Leone et al., 2015; Thaiss et al., 2014; Tofani, Clarke, et al., 2025), the time when participants collected their stool samples were similarly distributed across the groups. Interindividual variation in bowel movement frequency and gut transit time (which we, unfortunately, did not measure) further complicate circadian standardization (Asnicar et al., 2021). Given these considerations, any circadian influences would most likely introduce random variability rather than a systematic bias. This means that the observed gut microbiota-stress associations are unlikely to reflect circadian confounds.

Our approach has several limitations. First, we inferred individual SCFA production capacity from 16S rRNA gene amplicon sequencing data rather than directly measuring SCFA concentrations. Nevertheless, Frolova et al. (2022) found that SCFA production capacity significantly correlated with SCFA levels *in vitro* (specifically butyrate), supporting proxy validity. The sequencing approach may also have affected the identification and quantification of potential SCFA producers (we identified a substantially higher abundance of potential propionate producers compared to prior work, warranting future validation with alternative sequencing approaches). While shotgun metagenomic sequencing would provide a more direct assessment of microbial metabolic potential, our aim here was to provide first evidence linking inferred SCFA production capacity to stress-related parameters using a cost-effective and widely used 16S approach. Second, we did not detect a significant relationship between the gut microbiota and post-stress recovery. While this could indeed reflect a true null effect, we might have been underpowered, given the moderate group sizes of the stress and control groups. However, prior studies reported robust microbiota-stress associations with similar sample sizes (Delgadillo et al., 2025; Kato-Kataoka et al., 2016; Schmidt et al., 2015). It is also possible that any effects on post-stress recovery were masked by our stress reminder (i.e., to ensure a substantial effect of the acute stress intervention, we had repeated part of the stress procedure after a delay, which likely affected the trajectory of post-stress recovery). Third, although the distribution of biological sexes did not differ significantly between groups, the subgroup sizes were relatively uneven (see **Supplemental Figure S4** for an overview of sex-stratified group averages of cortisol/subjective stress trajectories). Details of the post-hoc power analysis still indicated adequate power to detect medium-to-large-sized effects (**Supplementary Results 6**). These effect sizes fall within the range typically reported in human studies linking the gut microbiota to behavioral and physiological variables (Nearing et al., 2022; Vujkovic-Cvijin et al., 2020). We recommend that future studies use larger and more evenly distributed participant samples and integrate relevant behavioral, microbial, and physiological markers of post-stress recovery and potential differences pertaining to biological sex.

To conclude, we investigated the role of the gut microbiota in modulating acute stress in healthy adults. Higher microbial diversity was associated with higher (both cortisol and subjective) stress reactivity after acute stress. Cortisol stress reactivity was also tied to the inferred capacity to produce key SCFAs, with higher butyrate and propionate production capacity revealing opposing relationships with stress parameters. These results are the first to highlight the link between gut microbial diversity, SCFA production capacity, and the acute stress response in healthy adults, underscoring the gut microbiota’s potential to modulate human psychophysiology in the aftermath of stress.

## 5. Acknowledgements

We thank Luise Graichen, Lars Keuter, Anna Schranz, Niklas Richter, and Magdalena Linder for help with study coordination and data collection. We also thank Gudrun Kohl, Jasmin Schwarz, and Joana Silva for DNA extraction and for processing the amplicon sequencing data, and especially Joana Silva for helpful comments on the manuscript. The amplicon sequencing data were processed using the Life Science Computer Cluster (LiSC) of the University of Vienna. This research was funded in part by the Austrian Science Fund (FWF) [10.55776/P34775], a NARSAD Young Investigator Grant of the Brain & Behavior Research Foundation (31532), and by the European Research Council (Starting Grant 101164099), awarded to I.C.W. P.A.G.F. was supported by a Marie Skłodowska-Curie Postdoctoral Fellowship from the European Commission (Grant number: 101107160). D.B. was supported by the FWF (10.55776/COE7, 10.55776/FG29). For open access purposes, the author has applied a CC BY public copyright license to any author-accepted manuscript version arising from this submission.

## 6. Author contributions

Conceptualization: T.K., I.C.W.; Data curation: T.K., I.C.W.; Formal analysis: T.K., I.C.W.; Funding acquisition: I.C.W.; Investigation: T.K., I.C.W.; Methodology: all authors; Project administration: T.K., I.C.W.; Resources: I.C.W.; Software: T.K., I.C.W.; Supervision: I.C.W.; Validation: T.K., I.C.W.; Visualization: T.K., I.C.W.; Writing – original draft: T.K., I.C.W.; Writing – reviewing and editing: all authors.

