## Supplementary Materials for "Gut microbial diversity and inferred capacity to produce short-chain fatty acids are tied to acute stress reactivity in healthy adults"

***This document includes:***

Supplementary Results 1-6
Supplementary Figures 1-4
Supplementary Tables 1-12
References

***General note on data & code availability, and results reporting***

The raw 16S rRNA gene amplicon sequencing data is available at the Sequence Read Archive (https://www.ncbi.nlm.nih.gov/sra) under the BioProject accession PRJNA1366128. All other raw data, analysis code, and the complete model outputs are openly available via the Open Science Framework (10.17605/OSF.IO/RAVUY). Due to the wealth of models computed, we restricted our manuscript to reporting results that pertain to the main research questions. The complete model outputs of all (non-/significant) results, as well as all results pertaining to additional exploratory analyses, are provided open access on the OSF.

### Supplementary Results

#### *Supplementary Results 1: Effect of stool quality (stress group)*

Methodology. To assess the potential effect of stool quality (as measured by the Bristol Stool Chart; Lewis & and Heaton, 1997), we repeated all RLMs (focused on the stress group only, as this was our main group-of-interest, $n_{stress}$= 35) that had included gut microbiota parameters and added stool quality as a covariate. Model formula: Dependent variable ~ gut microbiota measure (e.g., gut microbial alpha diversity, SCFA-producing taxa) + biological sex + trait anxiety + perceived stress + stool quality + baseline cortisol ($T_2$) (the latter was only included for cortisol stress reactivity and post-stress recovery).

Results – Gut microbial alpha diversity. Results showed that the association between gut microbial alpha diversity (Shannon Index, Inverse Simpson Index, observed number of ASVs) and cortisol (adjusted variance-based $R^2$ ranging between 0.224-0.239) or subjective (adjusted variance-based $R^2$ ranging between 0.165-0.392) stress reactivity remained significant after accounting for stool quality across participants in the stress group (all $p < 0.05$, statistical details for these and all following results are provided via the OSF, 10.17605/OSF.IO/RAVUY).

Results – Inferred SCFA production capacity. Results showed that the association between the inferred SCFA production capacity (butyrate, propionate) and cortisol stress reactivity remained largely unchanged. For butyrate, the effect estimate remained similar to the original model ($b$ = 0.0038, $SE$ = 0.0019, $z$ = 1.98; $p$ = 0.048; adjusted variance-based $R^2$ = 0.14). For propionate, the association remained significant as well ($b$ = -0.0039, $SE$ = 0.0019, z = -2.03; $p$ = 0.042; adjusted variance-based $R^2$ = 0.15).

Conclusions. Overall, stool quality did not emerge as a significant predictor in any of the models (gut microbial alpha diversity, inferred SCFA production capacity), and including stool quality did not increase the variance explained when compared to our original models. Taken together, these results demonstrate that stool quality does not account for the reported link between the gut microbiota and stress-related parameters.

#### *Supplementary Results 2: Higher cortisol stress reactivity in biological males (stress group)*

Methodology & results. We conducted several robustness checks, all of which are suitable for modest and unbalanced samples (Delacre et al., 2017; Good, 2005; Mann & Whitney, 1947), and evaluated whether the higher cortisol stress reactivity in biological males was driven by the unequal distribution of biological males/females in the stress group. For these analyses, we utilized the same input data (i.e., with identical scaling and centering settings) as for the RLMs in the main analysis. First, a Welch's $t$-test confirmed a significant sex difference in cortisol reactivity ($t$ = -2.338; df = 15.702; $p$ = 0.033), with biological males showing higher mean cortisol reactivity than females in the stress group (male: 0.00874; female: -0.00058). The effect size was substantial (Hedges' $g$ = -0.88, 95% CI [-1.65, -0.11]) . Second, we performed non-parametric testing using a Mann-Whitney U-test. Results once more showed that biological males had a higher cortisol stress reactivity than biological females ($W$ = 62; $r$ = -0.39, $p$ = 0.022). Third, we performed non-parametric testing using a permutation test (10,000 iterations). Results supported significantly higher cortisol stress reactivity in biological males compared to females ($p$ = 0.023).

Conclusions. In summary, the additional analyses suggest that biological males (in the stress group) showed generally higher cortisol stress reactivity than biological females, and that this effect was not caused by the lower number of biological males in the group. However, we fully acknowledge that the distribution of biological sexes in our sample is not ideal (modest overall sample size, uneven subgroups), and we now mention this as an important limitation in our Discussion section.

***Supplementary Results 3: Effects of (cortisol and subjective) stress reactivity, stratified by biological sex***

Methodology. We addressed potential sex differences in cortisol dynamics and/or subjectively experienced stress in the stress group ($n_{\text{stress}} = 35$, $n_{\text{females}} = 25$, $n_{\text{males}} = 10$). Specifically, we computed robust linear models (RLMs) in which cortisol/subjective stress reactivity were predicted by the gut microbial parameters (gut microbial alpha diversity, inferred SCFA production capacity) stratified by biological sex.

Results – Cortisol stress reactivity. In biological females, all three gut microbial alpha diversity metrics (Shannon Index, Inverse Simpson Index, observed number of ASVs) showed significantly positive associations with cortisol stress reactivity (all $p < 0.05$; adjusted variance-based $R^2 = 0.197$-$0.244$). In biological males, none of the gut microbial alpha diversity metrics reached significance (all $p > 0.05$). There was no significant effect of inferred butyrate production capacity on cortisol stress reactivity in women ($p = 0.446$), but a significantly negative effect of inferred propionate production capacity ($p = 0.037$; adjusted variance-based $R^2 = 0.185$). Although inferred butyrate production capacity had a significant effect on cortisol stress reactivity in men ($p < 0.001$), the adjusted variance-based $R^2$ was strongly negative, suggesting unstable model estimates (no significant effect of propionate, $p = 0.765$).

Results – Subjective stress reactivity. In biological females, gut microbial alpha diversity (Shannon Index: $p = 0.011$, Inverse Simpson Index: $p = 0.054$, observed number of ASVs: $p = 0.092$, adjusted variance-based $R^2 = -0.034$ to $-0.094$) partly showed significantly positive associations with subjective stress reactivity. There were no significant associations in biological males (all $p > 0.05$). There was no significant effect of inferred SCFA-production capacity on subjective stress reactivity in women (all $p > 0.05$), but a significant effect of inferred propionate-production capacity in men ($p = 0.045$; adjusted variance-based $R^2 = 0.240$).

Conclusions. Importantly, in both biological females and males, the signs of the coefficients for all gut microbial predictors (gut microbial alpha diversity, inferred SCFA production capacity) were consistent with the direction of the main results. Taken together, sex-stratified analyses did not alter the interpretation of our main findings. Associations observed in the full sample reappeared primarily in biological females but were weaker, while effects in biological males were highly variable and statistically unreliable due to limited power (which is not surprising, given the low number of biological males in the stress group). The models that included the full sample thus provided the most stable and interpretable estimates.

***Supplementary Results 4: Analysis of cortisol dynamics using area under the curve (AUC)***

Background & methodology. Similar to the summary measures of cortisol reactivity and post-stress recovery, the area under the curve (AUC) represents another summary measure that captures cortisol dynamics after acute stress (Pruessner et al., 2003). We modelled cortisol responses as the area under the curve (AUC) with respect to increase (AUCi) and ground (AUCg; collapsing across cortisol stress reactivity and post-stress recovery timepoints). We computed AUGg (i.e., total cortisol output over the sampling period) and AUCi (i.e., cumulative increase above baseline) using the trapezoidal formulas of Pruessner and colleagues (Pruessner et al., 2003). Identical to our main analysis, the AUCg and AUCi values were log-transformed and assessed using a robust regression framework. Baseline cortisol ($T_2$) was not included as a covariate in AUCi models, since it was already accounted for in the formula:

$$\mathrm{AUCg} = \frac{\mathrm{T}_2 + T_3}{2} * Time\ in\ minutes(T_3 - T_2) + \frac{\mathrm{T}_3 + T_4}{2} * Time\ in\ minutes(T_4 - T_3) + \frac{\mathrm{T}_5 + T_4}{2} * Time\ in\ minutes(T_5 - T_4)$$

$$\mathrm{AUCi} = AUCg - (\mathrm{T}_2 * Time\ in\ minutes((T_3 - T_2) + (T_4 - T_3) + (T_5 - T_4))$$

Log-transformations:

$$logAUCg = \log(AUCg + 1)$$

$$logAUCi = sign(AUCi) * \log(|AUCi| + 1)$$

In line with our calculations regarding (cortisol or subjective) stress reactivity, we employed the following robust linear model (RLM) for the AUC calculations (i.e., AUC predicted as part of a group interaction model, and models focusing on the stress/control groups separately):

Dependent variable ~ gut microbiota measure (e.g., gut microbial alpha diversity, abundance of SCFA-producing taxa) + biological sex + trait anxiety + perceived stress + baseline cortisol ($T_2$; note that the latter was only included for AUCg)

<u>Results – Gut microbial alpha diversity.</u> The AUCi models replicated the main findings, showing significant gut microbial alpha diversity (Shannon Index, Inverse Simpson Index, observed number of ASVs) × group interactions for both the Shannon and Inverse Simpson models (separate RLMs, $N_{\mathrm{total}} = 74$, $n_{\mathrm{stress}} = 35$, $n_{\mathrm{control}} = 39$; both $p < 0.05$), but not for the observed number of ASVs model ($p = 0.125$). Overall, this indicated that the association between gut microbial alpha diversity and AUCi was stronger and more positive in the stress group compared to controls (statistical details for these and all following results are provided via the OSF, 10.17605/OSF.IO/RAVUY). Similarly, for the AUCg models, we found a significant gut microbial alpha diversity × group interaction for all three alpha diversity metrics (separate RLMs, $N_{\mathrm{total}} = 74$, $n_{\mathrm{stress}} = 35$, $n_{\mathrm{control}} = 39$; all $p < 0.05$). In other words, higher gut microbial alpha diversity was more strongly associated with higher AUCg values across those participants who had undergone the acute stress intervention. When focusing exclusively on the stress group (separate RLMs, $n_{stress} = 35$), higher gut microbial alpha diversity was significantly associated with higher AUCi and AUCg values (all $p < 0.05$). There were no significant effects in the control group (separate RLMs, $n_{\mathrm{control}} = 39$, all $p > 0.05$).

<u>Results – Inferred SCFA production capacity.</u> Results from the AUCi models (separate RLMs, $N_{\mathrm{total}} = 74$, $n_{\mathrm{stress}} = 35$, $n_{\mathrm{control}} = 39$) revealed significant SCFA (butyrate, propionate) × group interactions for propionate ($p = 0.025$), but not butyrate ($p = 0.107$; statistical details for these and all following results are provided via the OSF, 10.17605/OSF.IO/RAVUY). The AUCg models revealed a significant SCFA × group interaction for butyrate ($p = 0.017$), but not propionate ($p = 0.068$). In other words, higher inferred SCFA production capacity was more strongly associated with lower AUCi (propionate) and higher AUCg (butyrate) values across those participants who had undergone the acute stress intervention. Within the stress group (separate RLMs, $n_{\mathrm{stress}} = 35$), a higher relative abundance of butyrate-producing bacteria was associated higher AUCg ($p = 0.042$, but not with AUCi, $p = 0.131$), whereas a higher relative

abundance of propionate-producing bacteria was significantly associated with lower AUCi ($p = 0.032$) and AUCg ($p = 0.035$). No significant effects were observed in the control group (separate RLMs, $n_{control} = 39$; all $p > 0.05$).

***Supplementary Results 5: Analysis of cortisol dynamics on a per-timepoint basis***

Background & Methodology. To complement the analyses of cortisol dynamics using summary measures such as cortisol stress reactivity or post-stress recovery (see our main analysis), and the area under the curve (**Supplementary Results 4**), we conducted additional linear mixed-effects models (LMMs) to characterize cortisol dynamics on a per-timepoint basis. For this, we utilized all available cortisol samples pertaining to the full sampling period as defined by the cortisol stress reactivity and post-stress recovery phases ($T_2$ to $T_5$) of both the stress and control groups ($N_{total} = 74$, $n_{stress}=35$, $n_{control}=39$). This allowed us to test whether associations with gut microbiota features (gut microbial alpha diversity, abundance of SCFA-producing taxa) were detectable across the entire cortisol trajectory, while accounting for repeated measures within individuals. In cases where the models were unable to converge, the following parameters were applied: lmerControl(optimizer = "bobyqa", optCtrl = list(maxfun = 100000)). The models were specified as follows:

logCortisol ~ time in minutes from stress onset * gut microbiota measure (gut microbial alpha diversity, abundance of SCFA-producing taxa) * group + biological sex + trait anxiety + perceived stress

Results – Gut microbial alpha diversity. Results confirmed that cortisol trajectories varied as a function of gut microbial alpha diversity (Shannon Index, Inverse Simpson Index, observed number of ASVs) and group (stress, control; separate LMMs, $N_{total} = 74$, $n_{stress} = 35$, $n_{control} = 39$). The significant time × group interactions (all $p < 0.01$) suggested that cortisol values were higher in the stress group compared to controls. Moreover, we found significant time × alpha diversity × group interactions (all $p < 0.05$) for all three metrics (statistical details for these and all following results are provided via the OSF, 10.17605/OSF.IO/RAVUY). This means that higher gut microbial alpha diversity was associated with higher cortisol values in the stress group after onset of acute stress (compared to controls) over time. Similarly, when focusing on the stress group alone (separate LMMs, $n_{stress} = 35$), higher values in all three alpha diversity metrics were significantly associated with higher cortisol values after onset of acute stress (all $p < 0.05$). There were no significant effects in the control group (separate LMMs, $n_{control} = 39$; all $p > 0.05$).

Results – Inferred SCFA production capacity. Cortisol trajectories varied as a function of inferred SCFA production capacity (butyrate, propionate) and group (stress, control; separate LMMs, $N_{total} = 74$, $n_{stress} = 35$, $n_{control} = 39$). For propionate, we observed a significant time × SCFA × group interaction ($p = 0.033$), whereas no significant effect was found for butyrate ($p = 0.172$; statistical details for these and all following results are provided via the OSF, 10.17605/OSF.IO/RAVUY). Thus, in line with our main results, a higher relative abundance of potential propionate-producing bacteria was associated with lower cortisol values in the stress group (compared to controls) over time. There were no significant effects when testing the stress (separate LMMs, $n_{stress} = 35$; all $p > 0.05$) or control groups separately (separate LMMs, $n_{control} = 39$; all $p > 0.05$).

***Supplementary Results 6: Post-hoc power analysis***

Background & Methodology. To highlight that our sample size was well-suited to detect associations between the gut microbiota and stress-related parameters, we conducted a post-hoc power analysis. We focused on two of our main models that tested the effect of 1) gut microbial alpha diversity (Shannon Index) and 2) inferred SCFA production capacity (butyrate) on cortisol stress reactivity in the stress group (separate RLMs, $n_{stress}$ = 35). We then utilized the effect sizes from these models and determined the proportion of variance explained using (adjusted) variance-based $R^2$. Because our main analyses used robust linear models (RLMs), classical $R^2$ is not defined for M-estimators, and to our best knowledge, no dedicated power analysis routines for RLMs exist. Therefore, we relied on the variance-based $R^2$ values derived from the RLMs as approximations to quantify effect magnitudes. Subsequently, we used the "pwr" package (version 1.3; Champely, 2020) to compute the achieved post-hoc power at $\alpha$ = 0.05 based on (i) the sample size of the stress group ($n_{stress}$ = 35), (ii) the number of predictors in the model, and (iii) the estimated effect size (variance-based $R^2$). Note that these values should be interpreted as approximations of detectable effect sizes for robust regressions, rather than as exact values.

Results – Gut microbial alpha diversity (Shannon Index). When testing the effect of gut microbial alpha diversity (Shannon Index) on cortisol stress reactivity, the RLM explained ~36% of the variance (variance-based $R^2$ = 0.355; adjusted variance-based $R^2$ = 0.243). This corresponds to medium-to-large sized Cohen's $f^2$ = 0.55 (variance-based $R^2$) and $f^2$ = 0.32 (adjusted variance-based $R^2$). With a sample size of $n_{stress}$ = 35 and five model predictors (Shannon Index, biological sex, trait anxiety, perceived stress, baseline cortisol $T_2$), the achieved post-hoc power was 0.89 (variance-based $R^2$) and 0.64 (adjusted variance-based $R^2$) at $\alpha$ = 0.05. Sensitivity analysis indicated that, with our sample size and model structure, achieving 80% power would require effects explaining ≥ 30.6% of the variance (our model explained ~36% of the variance and exceeded this threshold).

Results – Inferred SCFA production capacity (butyrate). When testing for the effect of inferred SCFA-production capacity (butyrate) on cortisol stress reactivity, the RLM explained ~29% of the variance (variance-based $R^2$ = 0.294; adjusted variance-based $R^2$ = 0.173). This corresponds to medium-to-large sized Cohen's $f^2$ = 0.42 (variance-based $R^2$) and $f^2$ = 0.21 (adjusted variance-based $R^2$). With a sample size of $n_{stress}$ = 35 and five model predictors (butyrate-production capacity, biological sex, trait anxiety, perceived stress, baseline cortisol $T_2$), the achieved post-hoc power was 0.77 (variance-based $R^2$) and 0.44 (adjusted variance-based $R^2$). As above, sensitivity analysis indicated that with our sample size and model structure, achieving 80% power would require effects explaining ≥ 30.6% of the variance (our model explained ~29% of the variance and fell slightly below this threshold).

To summarize, we fully agree that replication in larger samples is essential. However, based on the post-hoc power analysis, we conclude that our study was adequately powered to detect medium-to-large-sized effects (but not small associations). Notably, this range of effect sizes aligns with those of other studies reporting an association between the gut microbiota, behavioral, and physiological variables in human participants (Nearing et al., 2022; Vujkovic-Cvijin et al., 2020).

### Supplementary Figures

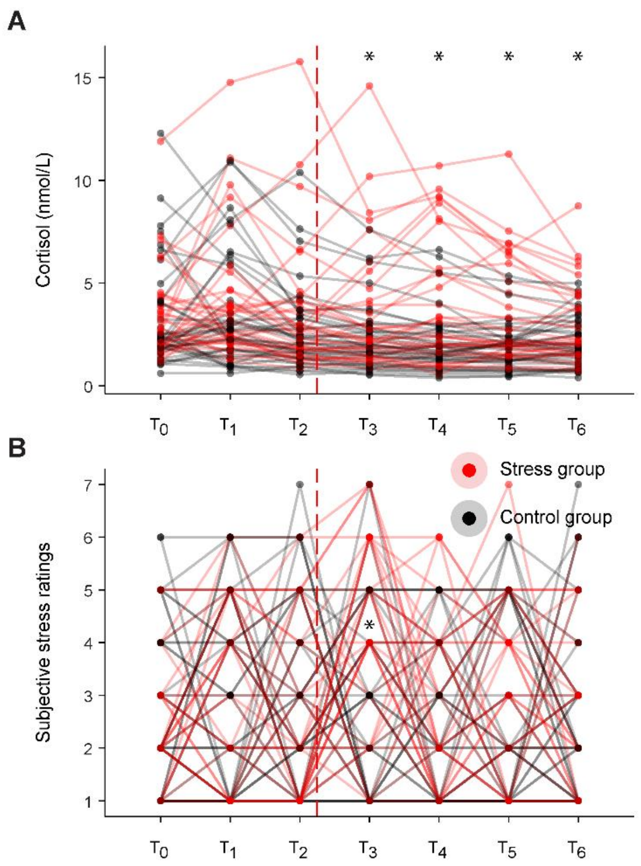


**Figure S1. Participant-specific trajectories of cortisol dynamics and subjectively experienced stress.** Individual trajectories of salivary cortisol concentrations and subjectively experienced stress over time. Timepoints (T) indicate the collection of salivary cortisol and ratings of subjectively experienced stress. Day 1 includes the $T_0$ measurement, and Day 2 comprises the remaining assessments ($T_1$–$T_6$). **(A)** Individual salivary cortisol concentrations (nmol/L; statistical comparisons were performed using the log-transformed values) for the stress ($n$ = 35) and control ($n$ = 39) groups across $T_0$–$T_6$. **(B)** Individual ratings of subjectively experienced stress across the same timepoints. Note that the connection between $T_0$ (Day 1) and $T_1$ (Day 2) does not imply a continuous within-day trajectory; it is included solely to indicate which $T_0$ value belongs to which participant. The dashed red line indicates the timing of the acute stress intervention (at $T_2$). * denotes significant group differences ($p < 0.05$).

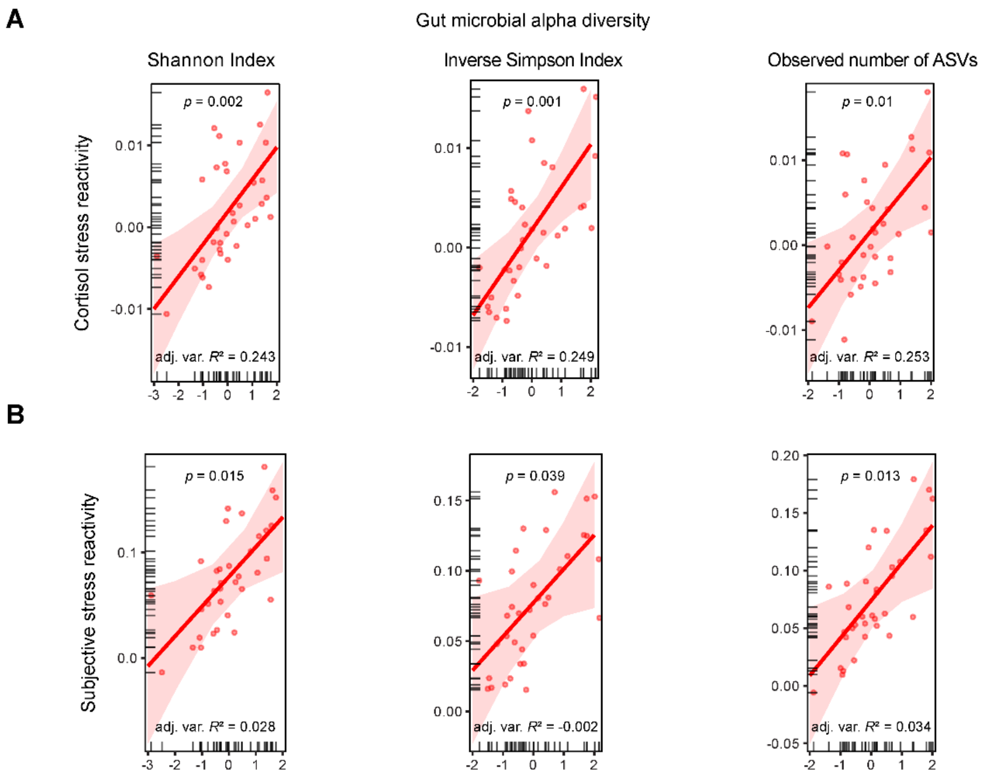


**Figure S2. Gut microbial alpha diversity is associated with cortisol and subjective stress reactivity in the stress group.** Partial effects from robust linear models (RLMs) predicting **(A)** cortisol stress reactivity (log-transformed concentrations, nmol/L) and **(B)** subjective stress reactivity based on gut microbial alpha diversity (Shannon Index, Inverse Simpson Index, observed number of ASVs) in the stress group. Data points represent participant-specific fitted values; tick marks oriented toward the inside of the plot area indicate the distribution of the observed data points along the predictor axes; shaded areas around the regression line represent the 95% confidence intervals; *p*-values indicate significant effects of gut microbial alpha diversity on cortisol or subjective stress reactivity; adjusted variance-based *R*2 values (shown as adj. var. *R*2) indicate overall model fit.

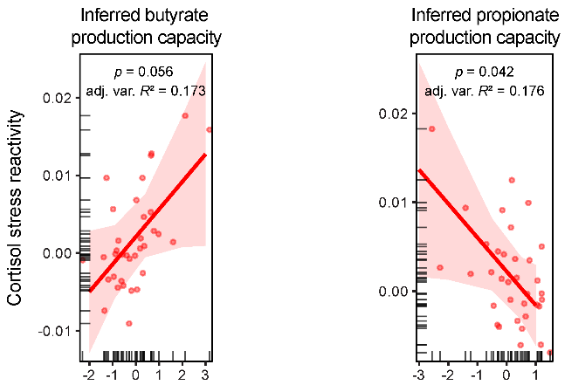


**Figure S3. Inferred gut microbial capacity to produce butyrate and propionate is associated with cortisol stress reactivity in the stress group.** Partial effects from robust linear models (RLMs) predicting cortisol stress reactivity (log-transformed concentrations, nmol/L) based on inferred SCFA production capacity for butyrate (left panel) and propionate (right panel) in the stress group. Data points represent participant-specific fitted values; tick marks oriented toward the inside of the plot area indicate the distribution of the observed data points along the predictor axes; shaded areas around the regression line represent the 95% confidence intervals; *p*-values indicate significant effects of inferred SCFA production capacity on cortisol stress reactivity; adjusted variance-based *R*2 values (shown as adj. var. *R*2) indicate overall model fit.

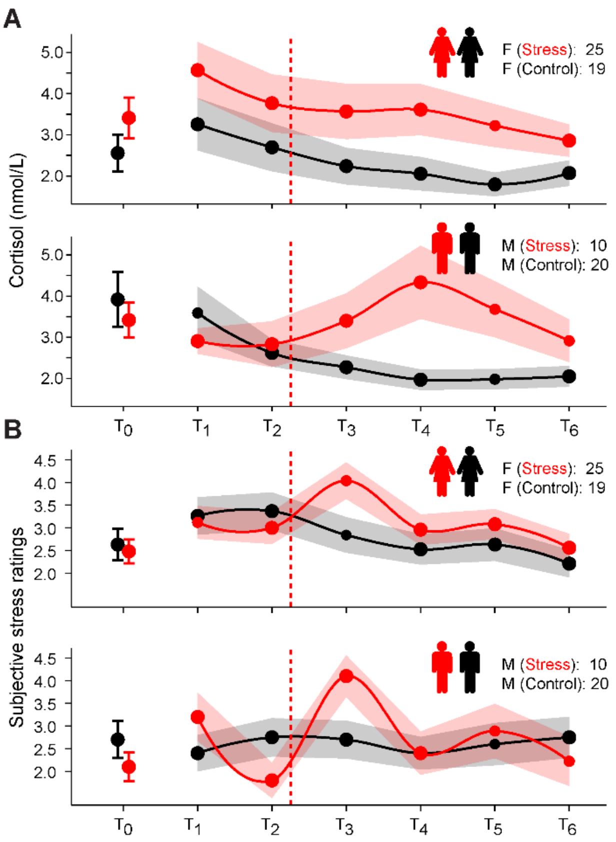


**Figure S4. Cortisol trajectories and subjectively experienced stress over time stratified by biological sex.** Trajectories of salivary cortisol concentrations and subjectively experienced stress over time, stratified by biological sex (females: $n_{stress}$ = 25, $n_{control}$ = 19; males: $n_{stress}$ = 10, $n_{control}$ = 20). Timepoints (T) indicate the collection of salivary cortisol and ratings of subjectively experienced stress. Day 1 includes the $T_0$ measurement, and Day 2 comprises the remaining assessments ($T_1$–$T_6$). **(A)** Salivary cortisol concentrations (nmol/L) for females (top panel) and males (bottom panel), and **(B)** subjective stress ratings for females (top panel) and males (bottom panel). Red lines indicate the stress group and black lines the control group for each biological sex. Continuous lines depict the respective group average, and shaded areas represent the standard error of the mean (SEM).

### Supplementary Tables

**Table S1. Cortisol dynamics and subjectively experienced stress over time.** Results from separate one-way ANOVAs testing for the effect of group (stress, control) on cortisol or subjective stress ratings at each sampling timepoint. Post-hoc *t*-tests for post-stress timepoints ($T_3$-$T_6$) were adjusted for multiple comparisons (using Bonferroni-Holm correction): * *p.adj* < .05.

| Timepoint | DFn | DFd | F | *p* | ges | *p.adj* |
|---|---|---|---|---|---|---|
| *Cortisol dynamics, main effect of group* | | | | | | |
| T0 (Day 1) | 1 | 72 | 0.59 | 0.445 | 0.008 | |
| T1 (Day 2 *ff*) | 1 | 72 | 2.154 | 0.147 | 0.029 | |
| T2 | 1 | 72 | 2.273 | 0.136 | 0.031 | |
| T3 | 1 | 72 | 5.867 | 0.018 | 0.075 | 0.036* |
| T4 | 1 | 72 | 11.218 | 0.001 | 0.135 | 0.004* |
| T5 | 1 | 72 | 12.02 | 0.001 | 0.143 | 0.004* |
| T6 | 1 | 72 | 5.216 | 0.025 | 0.068 | 0.036* |
| *Subjective stress ratings, main effect of group* | | | | | | |
| T0 (Day 1) | 1 | 72 | 0.748 | 0.390 | 0.010 | |
| T1 (Day 2 *ff*) | 1 | 72 | 0.600 | 0.441 | 0.008 | |
| T2 | 1 | 72 | 0.898 | 0.347 | 0.012 | |
| T3 | 1 | 72 | 9.382 | 0.003 | 0.115 | 0.012* |
| T4 | 1 | 72 | 0.858 | 0.357 | 0.012 | 0.966 |
| T5 | 1 | 71 | 0.995 | 0.322 | 0.014 | 0.966 |
| T6 | 1 | 71 | 0.002 | 0.966 | 0.00003 | 0.966 |

**Table S2. Gut microbial alpha diversity predicting cortisol stress reactivity across groups (interaction with biological sex, post-hoc RLMs).** Results from robust linear regression models (RLMs) predicting cortisol stress reactivity across groups based on gut microbial alpha diversity (Shannon Index, Inverse Simpson Index, observed number of ASVs), baseline cortisol, biological sex, and psychological variables (trait anxiety, perceived stress). Significance levels (Sig.): * $p < .05$, ** $p < .005$, *** $p < .0001$. Model formulas: Cortisol stress reactivity ~ biological sex * group + trait anxiety + perceived stress + alpha diversity metric * group + baseline cortisol ($T_2$).

| Predictor | *b* | SE | *z* | *p* | Sig. |
|---|---|---|---|---|---|
| *RLM (Shannon Index) predicting cortisol stress reactivity* | | | | | |
| (Intercept) | -0.005 | 0.001 | -3.35 | 0.001 | *** |
| Biological sex | 0.0004 | 0.002 | 0.193 | 0.847 | |
| Trait anxiety | 0.004 | 0.001 | 3.124 | 0.002 | ** |
| Perceived stress | -0.003 | 0.001 | -2.273 | 0.023 | * |
| Shannon | 0.0001 | 0.001 | 0.073 | 0.942 | |
| Stress group | 0.004 | 0.002 | 1.996 | 0.046 | * |
| Baseline cortisol ($T_2$) | -0.002 | 0.001 | -2.423 | 0.015 | * |
| Shannon:Stress group | 0.004 | 0.001 | 2.622 | 0.009 | ** |
| Biological sex:Stress group | 0.01 | 0.003 | 3.255 | 0.001 | ** |
| *RLM (Inverse Simpson Index) predicting cortisol stress reactivity* | | | | | |
| (Intercept) | -0.005 | 0.001 | -3.43 | 0.001 | *** |
| Biological sex | 0.0003 | 0.002 | 0.153 | 0.879 | |
| Trait anxiety | 0.004 | 0.001 | 3.646 | 0.0003 | *** |
| Perceived stress | -0.003 | 0.001 | -2.685 | 0.007 | ** |
| Inverse Simpson | -0.0002 | 0.001 | -0.152 | 0.879 | |
| Stress group | 0.004 | 0.002 | 2.159 | 0.031 | * |
| Baseline cortisol ($T_2$) | -0.002 | 0.001 | -2.168 | 0.03 | * |
| Inverse Simpson:Stress group | 0.004 | 0.001 | 2.984 | 0.003 | ** |
| Biological sex:Stress group | 0.008 | 0.003 | 2.824 | 0.005 | ** |
| *RLM (observed number of ASVs) predicting cortisol stress reactivity* | | | | | |
| (Intercept) | -0.005 | 0.001 | -3.205 | 0.001 | ** |
| Biological sex | 0.0004 | 0.002 | 0.22 | 0.826 | |
| Trait anxiety | 0.004 | 0.001 | 3.19 | 0.001 | ** |
| Perceived stress | -0.003 | 0.001 | -2.268 | 0.023 | * |
| Observed | 0.0001 | 0.001 | 0.061 | 0.951 | |
| Stress group | 0.004 | 0.002 | 1.917 | 0.055 | |

| | | | | | |
|---|---|---|---|---|---|
| Baseline cortisol ($T_2$) | -0.002 | 0.001 | -2.515 | 0.012 | * |
| Observed:Stress group | 0.004 | 0.002 | 2.653 | 0.008 | ** |
| Biological sex:Stress group | 0.009 | 0.003 | 2.74 | 0.006 | ** |

**Table S3. Gut microbial alpha diversity predicting cortisol stress reactivity in the stress group.** Results from robust linear regression models (RLMs) predicting cortisol stress reactivity in the stress group based on gut microbial alpha diversity (Shannon Index, Inverse Simpson Index, observed number of ASVs), baseline cortisol, biological sex, and psychological variables (trait anxiety, perceived stress). Significance levels (Sig.): * $p < .05$, ** $p < .005$, *** $p < .0001$. Model formulas: Cortisol stress reactivity ~ biological sex + trait anxiety + perceived stress + alpha diversity metric * group + baseline cortisol ($T_2$). Complete (non-significant) results for the control group are available via the OSF (10.17605/OSF.IO/RAVUY).

| Predictor | *b* | SE | *z* | *p* | Sig. |
|---|---|---|---|---|---|
| *RLM (Shannon Index) predicting cortisol stress reactivity* | | | | | |
| (Intercept) | -0.001 | 0.002 | -0.422 | 0.673 | |
| Biological sex | 0.010 | 0.003 | 3.413 | 0.001 | *** |
| Trait anxiety | 0.004 | 0.002 | 1.598 | 0.110 | |
| Perceived stress | -0.003 | 0.002 | -1.392 | 0.164 | |
| Shannon | 0.004 | 0.001 | 3.148 | 0.002 | ** |
| Baseline cortisol ($T_2$) | -0.001 | 0.001 | -0.540 | 0.589 | |
| *RLM (Inverse Simpson Index) predicting cortisol stress reactivity* | | | | | |
| (Intercept) | -0.0003 | 0.002 | -0.212 | 0.832 | |
| Biological sex | 0.009 | 0.003 | 2.973 | 0.003 | ** |
| Trait anxiety | 0.005 | 0.002 | 2.064 | 0.039 | * |
| Perceived stress | -0.004 | 0.002 | -1.795 | 0.073 | |
| Inverse Simpson | 0.004 | 0.001 | 3.450 | 0.001 | *** |
| Baseline cortisol ($T_2$) | -0.0002 | 0.001 | -0.118 | 0.906 | |
| *RLM (observed number of ASVs) predicting cortisol stress reactivity* | | | | | |
| (Intercept) | -0.001 | 0.002 | -0.304 | 0.762 | |
| Biological sex | 0.009 | 0.004 | 2.569 | 0.010 | * |
| Trait anxiety | 0.006 | 0.003 | 2.005 | 0.045 | * |
| Perceived stress | -0.004 | 0.003 | -1.556 | 0.120 | |
| Observed | 0.004 | 0.002 | 2.583 | 0.010 | ** |
| Baseline cortisol ($T_2$) | -0.002 | 0.002 | -0.961 | 0.337 | |

**Table S4. Gut microbial alpha diversity predicting post-stress recovery (cortisol) across groups.** Results from robust linear regression models (RLMs) predicting (cortisol) post-stress recovery across groups based on gut microbial alpha diversity (Shannon Index, Inverse Simpson Index, observed number of ASVs), baseline cortisol, biological sex, and psychological variables (trait anxiety, perceived stress). Significance levels (Sig.): * $p < .05$, ** $p < .005$, *** $p < .0001$. Model formulas: Cortisol post-stress recovery ~ biological sex + trait anxiety + perceived stress + alpha diversity metric * group + baseline cortisol ($T_2$).

| Predictor | *b* | SE | *z* | *p* | Sig. |
|---|---|---|---|---|---|
| *RLM (Shannon Index) predicting (cortisol) post-stress recovery* | | | | | |
| (Intercept) | -0.003 | 0.001 | -3.697 | 0.0002 | *** |
| Biological sex | -0.001 | 0.001 | -0.684 | 0.494 | |
| Trait anxiety | -0.002 | 0.001 | -2.34 | 0.019 | * |
| Perceived Stress | 0.001 | 0.001 | 1.177 | 0.239 | |
| Shannon | -0.001 | 0.001 | -0.897 | 0.37 | |
| Stress group | 0.0002 | 0.001 | 0.144 | 0.886 | |
| Baseline cortisol ($T_2$) | -0.002 | 0.001 | -4.279 | 0.00002 | *** |
| Shannon:Stress group | 0.001 | 0.001 | 0.548 | 0.584 | |
| *RLM (Inverse Simpson Index) predicting (cortisol) post-stress recovery* | | | | | |
| (Intercept) | -0.003 | 0.001 | -3.757 | 0.0002 | *** |
| Biological sex | -0.001 | 0.001 | -0.682 | 0.495 | |
| Trait anxiety | -0.002 | 0.001 | -2.15 | 0.032 | * |
| Perceived Stress | 0.001 | 0.001 | 0.981 | 0.326 | |
| Inverse Simpson | -0.001 | 0.001 | -1.301 | 0.193 | |
| Stress group | 0.0003 | 0.001 | 0.294 | 0.769 | |
| Baseline cortisol ($T_2$) | -0.002 | 0.001 | -4.199 | 0.00003 | *** |
| Inverse Simpson:Stress group | 0.001 | 0.001 | 1.327 | 0.185 | |
| *RLM (observed number of ASVs) predicting (cortisol) post-stress recovery* | | | | | |
| (Intercept) | -0.003 | 0.001 | -3.554 | 0.0004 | *** |
| Biological sex | -0.001 | 0.001 | -0.699 | 0.485 | |
| Trait anxiety | -0.002 | 0.001 | -2.433 | 0.015 | * |
| Perceived Stress | 0.001 | 0.001 | 1.267 | 0.205 | |
| Observed | -0.001 | 0.001 | -0.731 | 0.465 | |
| Stress group | 0.0002 | 0.001 | 0.142 | 0.887 | |
| Baseline cortisol ($T_2$) | -0.002 | 0.001 | -4.184 | 0.00003 | *** |
| Observed:Stress group | -0.0002 | 0.001 | -0.178 | 0.859 | |

**Table S5. Gut microbial alpha diversity predicting post-stress recovery (cortisol) in the stress group.** Results from robust linear regression models (RLMs) predicting (cortisol) post-stress recovery in the stress group based on gut microbial alpha diversity (Shannon Index, Inverse Simpson Index, observed number of ASVs), baseline cortisol, biological sex, and psychological variables (trait anxiety, perceived stress). Significance levels (Sig.): * $p < .05$, ** $p < .005$, *** $p < .0001$. Model formulas: Cortisol post-stress recovery ~ biological sex + trait anxiety + perceived stress + alpha diversity metric + baseline cortisol ($T_2$). Complete (non-significant) results for the control group are available via the OSF (10.17605/OSF.IO/RAVUY).

| Predictor | *b* | SE | *z* | *p* | Sig. |
|---|---|---|---|---|---|
| *RLM (Shannon Index) predicting (cortisol) post-stress recovery* | | | | | |
| (Intercept) | -0.003 | 0.001 | -2.706 | 0.007 | ** |
| Biological sex | -0.003 | 0.002 | -1.463 | 0.144 | |
| Trait anxiety | -0.003 | 0.001 | -1.886 | 0.059 | |
| Perceived Stress | 0.002 | 0.001 | 1.507 | 0.132 | |
| Shannon | -0.0004 | 0.001 | -0.577 | 0.564 | |
| Baseline cortisol ($T_2$) | -0.003 | 0.001 | -3.250 | 0.001 | ** |
| *RLM (Inverse Simpson Index) predicting (cortisol) post-stress recovery* | | | | | |
| (Intercept) | -0.003 | 0.001 | -2.765 | 0.006 | ** |
| Biological sex | -0.002 | 0.002 | -1.365 | 0.172 | |
| Trait anxiety | -0.002 | 0.001 | -1.807 | 0.071 | |
| Perceived Stress | 0.002 | 0.001 | 1.438 | 0.15 | |
| Inverse Simpson | 0.00001 | 0.001 | 0.015 | 0.988 | |
| Baseline cortisol ($T_2$) | -0.003 | 0.001 | -3.17 | 0.002 | ** |
| *RLM (observed number of ASVs) predicting (cortisol) post-stress recovery* | | | | | |
| (Intercept) | -0.003 | 0.001 | -2.458 | 0.014 | * |
| Biological sex | -0.003 | 0.002 | -1.467 | 0.142 | |
| Trait anxiety | -0.003 | 0.001 | -2.035 | 0.042 | * |
| Perceived Stress | 0.002 | 0.001 | 1.684 | 0.092 | |
| Observed | -0.001 | 0.001 | -1.141 | 0.254 | |
| Baseline cortisol ($T_2$) | -0.003 | 0.001 | -3.18 | 0.001 | ** |

**Table S6. Gut microbial alpha diversity predicting subjective stress reactivity in the stress group.** Results from robust linear regression models (RLMs) predicting subjective stress reactivity in the stress group based on gut microbial alpha diversity (Shannon Index, Inverse Simpson Index, observed number of ASVs), baseline cortisol, biological sex, and psychological variables (trait anxiety, perceived stress). Significance levels (Sig.): * $p < .05$, ** $p < .005$, *** $p < .0001$. Model formulas: Subjective stress reactivity ~ biological sex + trait anxiety + perceived stress + alpha diversity metric * group + baseline cortisol ($T_2$). Complete (non-significant) results for the control group are available via the OSF (10.17605/OSF.IO/RAVUY).

| Predictor | *b* | SE | *z* | *p* | Sig. |
|---|---|---|---|---|---|
| *RLM (Shannon Index) predicting subjective stress reactivity* | | | | | |
| (Intercept) | 0.059 | 0.015 | 3.879 | 0.0001 | *** |
| Biological sex | 0.057 | 0.028 | 2.001 | 0.045 | * |
| Trait anxiety | -0.037 | 0.022 | -1.663 | 0.096 | |
| Perceived Stress | 0.023 | 0.021 | 1.106 | 0.269 | |
| Shannon | 0.028 | 0.012 | 2.421 | 0.015 | * |
| *RLM (Inverse Simpson Index) predicting subjective stress reactivity* | | | | | |
| (Intercept) | 0.062 | 0.015 | 4.047 | 0.0001 | *** |
| Biological sex | 0.05 | 0.028 | 1.797 | 0.072 | |
| Trait anxiety | -0.036 | 0.022 | -1.652 | 0.098 | |
| Perceived Stress | 0.023 | 0.021 | 1.096 | 0.273 | |
| Inverse Simpson | 0.024 | 0.012 | 2.068 | 0.039 | * |
| *RLM (observed number of ASVs) predicting subjective stress reactivity* | | | | | |
| (Intercept) | 0.057 | 0.015 | 3.813 | 0.0001 | *** |
| Biological sex | 0.053 | 0.028 | 1.925 | 0.054 | |
| Trait anxiety | -0.032 | 0.022 | -1.456 | 0.145 | |
| Perceived Stress | 0.022 | 0.021 | 1.065 | 0.287 | |
| Observed | 0.033 | 0.013 | 2.487 | 0.013 | * |

**Table S7. Gut microbial alpha diversity predicting post-stress recovery (subjective ratings) in the stress group.** Results from robust linear regression models (RLMs) predicting (subjective) post-stress recovery in the stress group based on gut microbial alpha diversity (Shannon Index, Inverse Simpson Index, observed number of ASVs), biological sex and psychological variables (trait anxiety, perceived stress). Significance levels (Sig.): * $p < .05$, ** $p < .005$, *** $p < .0001$. Model formulas: Subjective post-stress recovery ~ biological sex + trait anxiety + perceived stress + alpha diversity metric. Complete (non-significant) results for the control group are available via the OSF (10.17605/OSF.IO/RAVUY).

| Predictor | *b* | SE | *z* | *p* | Sig. |
|---|---|---|---|---|---|
| *RLM (Shannon Index) predicting subjective post-stress recovery* | | | | | |
| (Intercept) | -0.038 | 0.016 | -2.351 | 0.019 | * |
| Biological sex | 0.0004 | 0.031 | 0.014 | 0.989 | |
| Trait anxiety | 0.034 | 0.024 | 1.388 | 0.165 | |
| Perceived Stress | -0.039 | 0.023 | -1.736 | 0.083 | |
| Shannon | -0.019 | 0.014 | -1.357 | 0.175 | |
| *RLM (Inverse Simpson Index) predicting subjective post-stress recovery* | | | | | |
| (Intercept) | -0.039 | 0.016 | -2.353 | 0.019 | * |
| Biological sex | -0.001 | 0.031 | -0.039 | 0.969 | |
| Trait anxiety | 0.037 | 0.024 | 1.513 | 0.13 | |
| Perceived Stress | -0.041 | 0.023 | -1.812 | 0.07 | |
| Inverse Simpson | -0.016 | 0.013 | -1.179 | 0.238 | |
| *RLM (observed number of ASVs) predicting subjective post-stress recovery* | | | | | |
| (Intercept) | -0.039 | 0.016 | -2.394 | 0.017 | * |
| Biological sex | -0.0001 | 0.031 | -0.005 | 0.996 | |
| Trait anxiety | 0.036 | 0.024 | 1.476 | 0.14 | |
| Perceived Stress | -0.041 | 0.022 | -1.846 | 0.065 | |
| Observed | -0.011 | 0.015 | -0.758 | 0.448 | |

**Table S8. SCFA production pathways and associated taxa adapted from Frolova et al. (2022), Table 1. Frontiers in Molecular Biosciences, licensed under CC-BY 4.0.** Overview of fermentation pathway variants for butyrate and propionate production in the human gut microbiota reference genomes, including the number of genomes and species per pathway and top associated taxa (table adapted from Frolova et al., 2022).

| Fermentation product | Pathway variants† | # Pathways | Top taxonomic groups |
|---|---|---|---|
| Butyrate | P1 | 1 | Clostridiaceae, Eubacteriaceae, Lachnospiraceae, Streptococcaceae |
| | P1+P2 | 2 | Clostridiales |
| | P1+P3 | 2 | Clostridiales, Acidaminococcaceae, Peptoniphilaceae |
| | P1+P4 | 2 | Clostridiales, Butyricimonas, Odoribacter |
| | P1+P2+P3 | 3 | Clostridium, Lachnoclostridium |
| | P1+P2+P4 | 13 | Porphyromonas, Clostridiales |
| | P2+P4 | 2 | Paraclostridium |
| | P1+P3+P4 | 3 | Fusobacterium |
| | P1+P2+P3+P4 | 4 | Fusobacterium |
| | P2 | 1 | Lachnoclostridium, Porphyromonas, Tannerella |
| | P4 | 1 | Alistipes, Micromonospora |
| Propionate | P1 | 1 | Bacteroidetes, Firmicutes, Akkermansia, Enterobacteria, Actinobacteria (Propionibacteria/Corynebacteria) |
| | P1+P2 | 2 | Peptostreptococcaceae |
| | P1+P3 | 2 | Enterobacteria, Veillonellaceae |
| | P1+P2+P3 | 3 | Intestinimonas |
| | P2 | 1 | Clostridiaceae, Lachnospiraceae, Peptostreptococcaceae |
| | P2+P3 | 2 | Clostridiaceae, Eubacteriaceae, Peptostreptococcaceae |
| | P3 | 1 | Diverse Clostridia, Peptoniphilus, Veillonella, Lactobacillus, Enterococcus, Listeria, Enterobacteria, Fusobacterium |

† Butyrate pathways: P1 (Acetyl-CoA), P2 (succinate), P3 (glutamate), P4 (lysine); Propionate pathways: P1 (succinate), P2 (lactate), P3 (propanediol).

**Table S9. Inferred gut microbial capacity to produce butyrate and propionate predicting cortisol stress reactivity in the stress group.** Results from robust linear regression models (RLMs) predicting cortisol stress reactivity in the stress group based on the inferred gut microbial capacity to produce SCFAs (butyrate, propionate), baseline cortisol, biological sex, and psychological variables (trait anxiety, perceived stress). Significance levels (Sig.): * $p < .05$, ** $p < .005$, *** $p < .0001$. Model formulas: Cortisol stress reactivity ~ biological sex + trait anxiety + perceived stress + SCFAs + baseline cortisol ($T_2$). Complete (non-significant) results for the control group are available via the OSF (10.17605/OSF.IO/RAVUY). The final model shows results predicting cortisol stress reactivity in the stress group based on both SCFAs combined (butyrate + propionate): Cortisol stress reactivity ~ biological sex + trait anxiety + perceived stress + butyrate + propionate + baseline cortisol ($T_2$).

| Predictor | *b* | SE | *z* | *p* | Sig. |
|---|---|---|---|---|---|
| *RLM (butyrate) predicting cortisol stress reactivity* | | | | | |
| (Intercept) | -0.00001 | 0.002 | -0.003 | 0.998 | |
| Biological sex | 0.009 | 0.004 | 2.112 | 0.035 | * |
| Trait anxiety | 0.004 | 0.003 | 1.112 | 0.266 | |
| Perceived Stress | -0.002 | 0.003 | -0.508 | 0.611 | |
| Butyrate | 0.004 | 0.002 | 1.913 | 0.056 | . |
| Baseline cortisol ($T_2$) | -0.002 | 0.002 | -1.123 | 0.261 | |
| *RLM (propionate) predicting cortisol stress reactivity* | | | | | |
| (Intercept) | 0.001 | 0.002 | 0.312 | 0.755 | |
| Biological sex | 0.007 | 0.004 | 1.834 | 0.067 | |
| Trait anxiety | 0.006 | 0.003 | 2.069 | 0.039 | * |
| Perceived Stress | -0.004 | 0.003 | -1.372 | 0.17 | |
| Propionate | -0.004 | 0.002 | -2.03 | 0.042 | * |
| Baseline cortisol (T2) | -0.001 | 0.002 | -0.638 | 0.524 | |
| *RLM (butyrate & propionate) predicting cortisol stress reactivity* | | | | | |
| (Intercept) | 0.001 | 0.002 | 0.785 | 0.433 | |
| Biological sex | 0.008 | 0.003 | 2.494 | 0.013 | * |
| Trait anxiety | 0.006 | 0.002 | 2.437 | 0.015 | * |
| Perceived Stress | -0.003 | 0.002 | -1.127 | 0.26 | |
| Butyrate | 0.004 | 0.001 | 2.876 | 0.004 | ** |
| Propionate | -0.005 | 0.002 | -2.979 | 0.003 | ** |
| Baseline cortisol (T2) | -0.002 | 0.001 | -1.129 | 0.259 | |

**Table S10. Inferred gut microbial capacity to produce butyrate and propionate predicting post-stress recovery (cortisol) in the stress group.** Results from robust linear regression models (RLMs) predicting (cortisol) post-stress recovery in the stress group based on the inferred gut microbial capacity to produce SCFAs (butyrate, propionate), baseline cortisol, biological sex, and psychological variables (trait anxiety, perceived stress). Significance levels (Sig.): * $p < .05$, ** $p < .005$, *** $p < .0001$. Model formulas: Cortisol post-stress recovery ~ biological sex + trait anxiety + perceived stress + SCFAs * group + baseline cortisol ($T_2$). Complete (non-significant) results for the control group are available via the OSF (10.17605/OSF.IO/RAVUY).

| Predictor | *b* | SE | *z* | *p* | Sig. |
|---|---|---|---|---|---|
| *RLM (butyrate) predicting cortisol post-stress recovery* | | | | | |
| (Intercept) | -0.003 | 0.001 | -2.649 | 0.008 | ** |
| Biological sex | -0.002 | 0.002 | -1.356 | 0.175 | |
| Trait anxiety | -0.003 | 0.001 | -1.837 | 0.066 | |
| Perceived Stress | 0.002 | 0.001 | 1.455 | 0.146 | |
| Butyrate | 0.0002 | 0.001 | 0.299 | 0.765 | |
| Baseline cortisol (T2) | -0.003 | 0.001 | -3.228 | 0.001 | ** |
| *RLM (propionate) predicting cortisol post-stress recovery* | | | | | |
| (Intercept) | -0.003 | 0.001 | -2.766 | 0.006 | ** |
| Biological sex | -0.002 | 0.002 | -1.373 | 0.17 | |
| Trait anxiety | -0.002 | 0.001 | -1.783 | 0.075 | |
| Perceived Stress | 0.002 | 0.001 | 1.444 | 0.149 | |
| Propionate | 0 | 0.001 | -0.002 | 0.998 | |
| Baseline cortisol (T2) | -0.003 | 0.001 | -3.323 | 0.001 | *** |

**Table S11. Inferred gut microbial capacity to produce butyrate and propionate predicting subjective stress reactivity in the stress group.** Results from robust linear regression models (RLMs) predicting subjective stress reactivity in the stress group based on the inferred gut microbial capacity to produce SCFAs (butyrate, propionate), biological sex and psychological variables (trait anxiety, perceived stress). Significance levels (Sig.): * $p < .05$, ** $p < .005$, *** $p < .0001$. Model formulas: Subjective stress reactivity ~ biological sex + trait anxiety + perceived stress + SCFAs. Complete (non-significant) results for the control group are available via the OSF (10.17605/OSF.IO/RAVUY).

| Predictor | *b* | SE | *z* | *p* | Sig. |
|---|---|---|---|---|---|
| *RLM (butyrate) predicting subjective stress reactivity* | | | | | |
| (Intercept) | 0.065 | 0.017 | 3.943 | 0.0001 | *** |
| Biological sex | 0.037 | 0.031 | 1.228 | 0.22 | |
| Trait anxiety | -0.043 | 0.024 | -1.841 | 0.066 | |
| Perceived Stress | 0.032 | 0.023 | 1.409 | 0.159 | |
| Butyrate | 0.009 | 0.014 | 0.692 | 0.489 | |
| *RLM (propionate) predicting subjective stress reactivity* | | | | | |
| (Intercept) | 0.066 | 0.016 | 4.224 | 0.00002 | *** |
| Biological sex | 0.039 | 0.029 | 1.366 | 0.172 | |
| Trait anxiety | -0.04 | 0.023 | -1.761 | 0.078 | |
| Perceived Stress | 0.029 | 0.021 | 1.369 | 0.171 | |
| Propionate | -0.01 | 0.014 | -0.713 | 0.476 | |

**Table S12. Inferred gut microbial capacity to produce butyrate and propionate predicting post-stress recovery (subjective ratings) in the stress group.** Results from robust linear regression models (RLMs) predicting (subjective) post-stress recovery in the stress group based on the inferred gut microbial capacity to produce SCFAs (butyrate, propionate), biological sex and psychological variables (trait anxiety, perceived stress). Significance levels (Sig.): * $p < .05$, ** $p < .005$, *** $p < .0001$. Model formulas: Subjective post-stress recovery ~ biological sex + trait anxiety + perceived stress + SCFAs. Complete (non-significant) results for the control group are available via the OSF (10.17605/OSF.IO/RAVUY).

| Predictor | *b* | SE | *z* | *p* | Sig. |
|---|---|---|---|---|---|
| *RLM (butyrate) predicting subjective post-stress recovery* | | | | | |
| (Intercept) | -0.041 | 0.015 | -2.648 | 0.008 | ** |
| Biological sex | 0.001 | 0.03 | 0.027 | 0.979 | |
| Trait anxiety | 0.042 | 0.022 | 1.871 | 0.061 | |
| Perceived Stress | -0.052 | 0.021 | -2.443 | 0.015 | * |
| Butyrate | -0.018 | 0.014 | -1.251 | 0.211 | |
| *RLM (propionate) predicting subjective post-stress recovery* | | | | | |
| (Intercept) | -0.039 | 0.017 | -2.309 | 0.021 | * |
| Biological sex | 0.002 | 0.032 | 0.07 | 0.944 | |
| Trait anxiety | 0.043 | 0.025 | 1.737 | 0.082 | |
| Perceived Stress | -0.044 | 0.023 | -1.941 | 0.052 | |
| Propionate | -0.009 | 0.016 | -0.558 | 0.577 | |
